# An amino acid transporter AAT1 plays a pivotal role in chloroquine resistance evolution in malaria parasites

**DOI:** 10.1101/2022.05.26.493611

**Authors:** Alfred Amambua-Ngwa, Katrina A. Button-Simons, Xue Li, Sudhir Kumar, Katelyn Vendrely Brenneman, Marco Ferrari, Lisa A. Checkley, Meseret T. Haile, Douglas A. Shoue, Marina McDew-White, Sarah M. Tindall, Ann Reyes, Elizabeth Delgado, Haley Dalhoff, James K. Larbalestier, Roberto Amato, Richard D. Pearson, Alexander B. Taylor, François H. Nosten, Umberto D’Alessandro, Dominic Kwiatkowski, Ian H. Cheeseman, Stefan H. I. Kappe, Simon V. Avery, David J. Conway, Ashley M. Vaughan, Michael T. Ferdig, Timothy J. C. Anderson

**Author notes:** These authors contributed equally to this work (Alfred Amambua-Ngwa, Katrina A. Button-Simons, Xue Li, Sudhir Kumar, Katelyn Vendrely Brenneman).

## Abstract

Malaria parasites break down host hemoglobin into peptides and amino acids in the digestive vacuole for export to the parasite cytoplasm for growth: interrupting this process is central to the mode of action of several antimalarial drugs. Mutations in the chloroquine (CQ) resistance transporter, *pfcrt*, located in the digestive vacuole membrane, confer CQ resistance in *Plasmodium falciparum*, but typically affect parasite fitness. However, the role of other parasite loci in the evolution of CQ resistance is unclear. Here we use a combination of population genomics, genetic crosses and gene editing to demonstrate that a second vacuolar transporter plays a key role in both resistance and compensatory evolution. Longitudinal genomic analyses of the Gambian parasites revealed temporal signatures of selection on an amino acid transporter (*pfaat1)* S258<u>L</u> variant which increased from 0-87% in frequency between 1984 and 2014 in parallel with *pfcrt1* K76<u>T</u>. Parasite genetic crosses then identified a chromosome 6 quantitative trait locus containing *pfaat1* that is selected by CQ treatment. Gene editing demonstrated that *pfaat1* S258<u>L</u> potentiates CQ-resistance but at a cost of reduced fitness, while *pfaat1* F313<u>S</u>, a common Southeast Asian polymorphism, reduces CQ-resistance while restoring fitness. Our analyses reveal hidden complexity in CQ-resistance evolution, suggesting that *pfaat1* may underlie regional differences in the dynamics of resistance evolution, and modulate parasite resistance or fitness by manipulating the balance between both amino acid and drug transport.

---

Drug resistance in microbial pathogens complicates control efforts. Therefore, understanding the genetic architecture and the complexity of resistance evolution is critical for resistance monitoring and the development of improved treatment strategies. In the case of malaria parasites, deployment of five classes of antimalarial drugs over the last half century have resulted in well-characterized hard and soft selective sweeps associated with drug resistance, with both worldwide dissemination and local origins of resistance driving drug resistance alleles across the range of *P. falciparum* ^1–3^. Chloroquine (CQ) monotherapy played a central role in an ambitious plan to eradicate malaria in the last century. Resistance to CQ was first observed in 1957 in Southeast Asia (SEA), and subsequently arrived and spread across Africa from the late 1970s, contributing to the end of this ambitious global eradication effort ^4^.

Resistance to CQ has been studied intensively. The CQ resistance transporter (*pfcrt*, chromosome [chr.] 7) was originally identified using a *P. falciparum* genetic cross conducted between a CQ-resistant SEA parasite and a CQ-sensitive South American parasite generated in a chimpanzee host ^5, 6^. Twenty years of intensive research revealed the mechanistic role of CRT in drug resistance ^7, 8^, its location in the digestive vacuole membrane, and its natural function transporting short peptides from the digestive vacuole into the cytoplasm ^9^. CQ kills parasites by interfering with hemoglobin digestion in the digestive vacuole, preventing conversion of heme, a toxic by-product of hemoglobin digestion, into inert hemozoin crystals. Parasites carrying CQ-resistance mutations at CRT transport CQ out of the food vacuole, away from the site of drug action ^7, 8^. The *pfcrt* K76<u>T</u> single nucleotide polymorphism (SNP) is widely used as a molecular marker for CQ resistance ^10^, while additional variants within *pfcrt* modulate levels of resistance to CQ ^11^ and other quinoline drugs ^12^, and determine associated fitness costs ^13^. While mutations in a second transporter located in the food vacuole membrane, the multidrug resistance transporter (*pfmdr1),* have been shown to modulate CQ resistance in some genetic backgrounds ^14^, the role of other genes in CQ resistance evolution remains unclear. In this work we sought to understand the contribution of additional parasite loci to CQ-resistance evolution using a combination of population genomics, experimental genetic crosses, and gene editing.

## Results

### Strong signatures of selection on pfaat1

Longitudinal population genomic data can provide compelling evidence of the evolution of drug resistance loci ^15^. We conducted a longitudinal whole genome sequence analysis of 600 *P. falciparum* genomes collected between 1984 and 2014 in The Gambia to examine signatures of selection under drug pressure (Supplementary Table 1). Following filtration using genotype missingness (<10%) and minor allele frequency (>2%), we retained 16,385 biallelic SNP loci from 321 isolates (1984 [134], 1990 [13], 2001 [34], 2008 [75], and 2014 [65]. The *pfcrt* K76<u>T</u> allele associated with CQ-resistance increased from 0% in 1984 to 70% in 2014. Intriguingly, there was also rapid allele frequency change on chr. 6: the strongest differentiation is seen at an S258<u>L</u> mutation in a putative amino acid transporter, *pfaat1* (PF3D7_0629500, chr. 6), which increased during the same time period from 0% to 87% (Fig. 1a). Assuming a generation time (mosquito to mosquito) of 6 months for malaria parasites, these changes were driven by selection coefficients of 0.17 for *pfaat1* S258<u>L</u>, and 0.11 for *pfcrt* K76<u>T</u> (Supplementary Fig. 1). Both *pfaat1* S258<u>L</u> and *pfcrt* K76<u>T</u> mutations were absent in 1984 samples, but present in 1990, suggesting that they arose and spread in a short time window. Both *pfaat1* and *pfcrt* showed similar temporal haplotype structures in The Gambia (Supplementary Fig. 2). These were characterized by almost complete replacement of well differentiated haplotypes at both loci between 1984 and 2014. During this period, we also observed major temporal changes in another known drug resistance locus (*pfdhfr*) (chr. 4) (Fig. 1b) ^16^. That these rapid changes in allele frequency occur at *pfcrt*, *pfaat1* and *pfdhfr*, but not elsewhere in the genome (Fig. 1b), provides unambiguous evidence for strong directional selection.

**Fig. 1.**
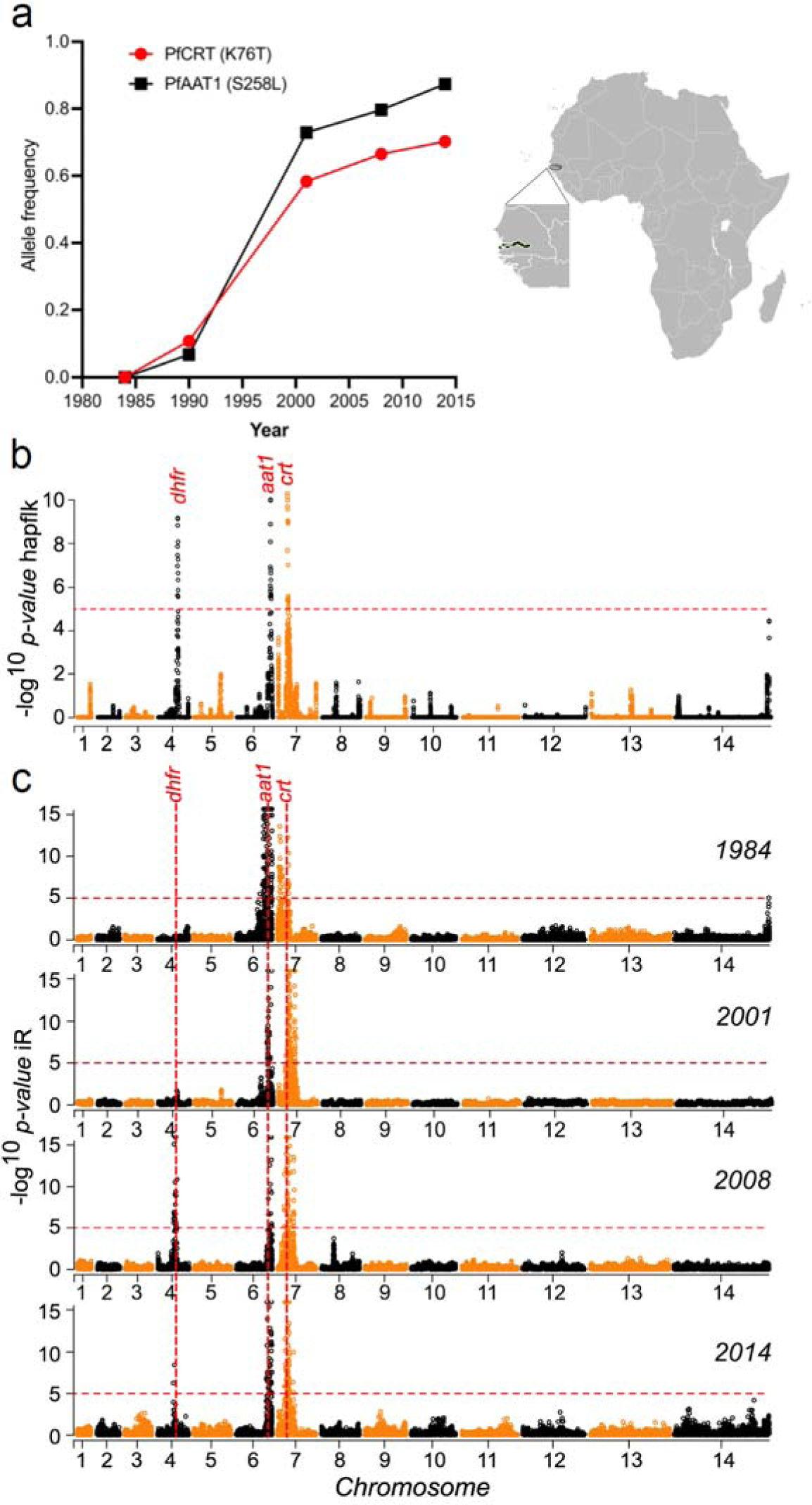
Rapid allele frequency change and strong signals of selection around *pfaat1* in The Gambia. (**a**) Temporal allele frequency change at SNPs coding for *pfaat1* S258<u>L</u> and *pfcrt* K76<u>T</u> between 1984 and 2014. The map and expanded West African region show the location of The Gambia. (**b**) Significance of haplotype differentiation across temporal populations of *P. falciparum* parasites determined using hapFLK. Significance thresholds at -log_10_(FDR-corrected *p*-value) = 5 are indicated with red dotted horizontal lines. Regions within the top 1% tail of FDR-corrected *p*-values are marked with gene symbols. The strongest signals genome-wide seen are around *pfcrt*, *pfaat1*, and *pfdhfr* (which is involved in pyrimethamine resistance). (**c**) Identity-by-descent (IBD), quantified with the isoRelate (iR) statistic, for temporal populations sampled from The Gambia. Consistently high peaks of IBD around *pfcrt* and *pfaat1* are seen for parasite populations in all years of sampling. The 1990 sample (n=13) is not shown in panel C.

Further evidence of strong selection on *pfaat1* and *pfcrt* came from the analysis of identity-by-descent (IBD) in Gambian parasite genomes. We saw the strongest signals of IBD in the genome around both *pfaat1* and *pfcrt* (Fig. 1c). These signals are dramatic, because there is minimal IBD elsewhere in the genome, with the exception of a strong signal centering on *pfdhfr* after 2008. Interestingly, the strong IBD is observed in all four temporal samples examined including 1984, prior to the spread of either *pfaat1* S258<u>L</u> or *pfcrt* K76<u>T</u>. However, only a single synonymous variant at *pfaat1* (I552I) and none of the CQ-resistant associated mutant variants in *pfcrt* were present at that time. CQ was the first line treatment across Africa from the 1950s. These results are consistent with the possibility of CQ-driven selective sweeps conferring low level CQ-resistance prior to 1984, perhaps targeting promotor regions of resistance-associated genes. *pfaat1* has also been selected in global locations: this is evident from prior population genomic analyses from Africa ^17^, SEA ^18^ and South America ^19^. Plots summarizing IBD in these regions are provided in Supplementary Fig. 3.

Patterns of linkage disequilibrium (LD) provide further evidence for functional linkage between *pfcrt* and *pfaat1*. The strongest genome-wide signal of interchromosomal LD was found between these two loci both in our Gambian data (Supplementary Fig. 4) and in samples from across Africa ^20^. LD between *pfaat1* and *pfcrt* was strongest in 2001, and then decayed in 2008 and 2014 (Supplementary Fig. 4&5), consistent with maintenance of LD during intensive CQ usage, and subsequent LD decay after CQ monotherapy was replaced by sulfadoxine-pyrimethamine + CQ combinations in 2004, and then with artemisinin combinations in 2008 ^16^.

Correlations in allele frequencies are expected between *pfcrt* and *pfaat1* if these loci are interacting or are co-selected. Frequencies of the CVIET haplotype for amino acids 72-76 in CRT are significantly correlated with allele frequencies of *pfaat1* S258<u>L</u> in West Africa (*R^2^* = 0.65, *p* = 0.0017) or across all African populations (*R^2^* = 0.44, *p* = 0.0021) (Supplementary Fig. 6). This analysis further strengthens the argument for co-evolution or epistasis between these two genes.

### Divergent selection on pfaat1 in Southeast Asia

We examined the haplotype structure of *pfaat1* from *P. falciparum* genomes (MalariaGEN release 6 ^21^) (Fig. 2, Supplementary Table 2). The *pfaat1* S258<u>L</u> SNP is at high frequency in SEA (58%) but is found on divergent flanking haplotypes suggesting an independent origin from the *pfaat1* S258<u>L</u> in The Gambia and elsewhere in Africa (Fig. 2 c & d, Supplementary Fig. 7). Hendon et al ^18^ reached the same conclusion for the chr. 6 region using an identity-by-descent analysis of parasites from global locations. Convergent evolution of *pfaat1* S258<u>L</u> provides further evidence for selection, and contrasts with *pfcrt* and *pfdhfr*, where resistance alleles that spread in Africa had an Asian origin ^1, 2^. *Pfaat1* evolution is more complex in SEA than elsewhere in the world. There are three additional common derived amino acid changes in SEA. *Pfaat1* F313<u>S</u> has spread close to fixation in SEA (total 96%, F_ST_ = 0.91 compared with African samples) paired with *pfaat1* S258<u>L</u> (55%), Q454<u>E</u> (15%), or K541<u>N</u> (22%). We speculate that these geographically localized *pfaat1* haplotypes have played an important role in CQ-resistance evolution in SEA and could also reflect geographic differences in the historical use of other quinoline drugs (mefloquine, quinine, piperaquine, lumefantrine) in this region ^22^.

**Fig. 2.**
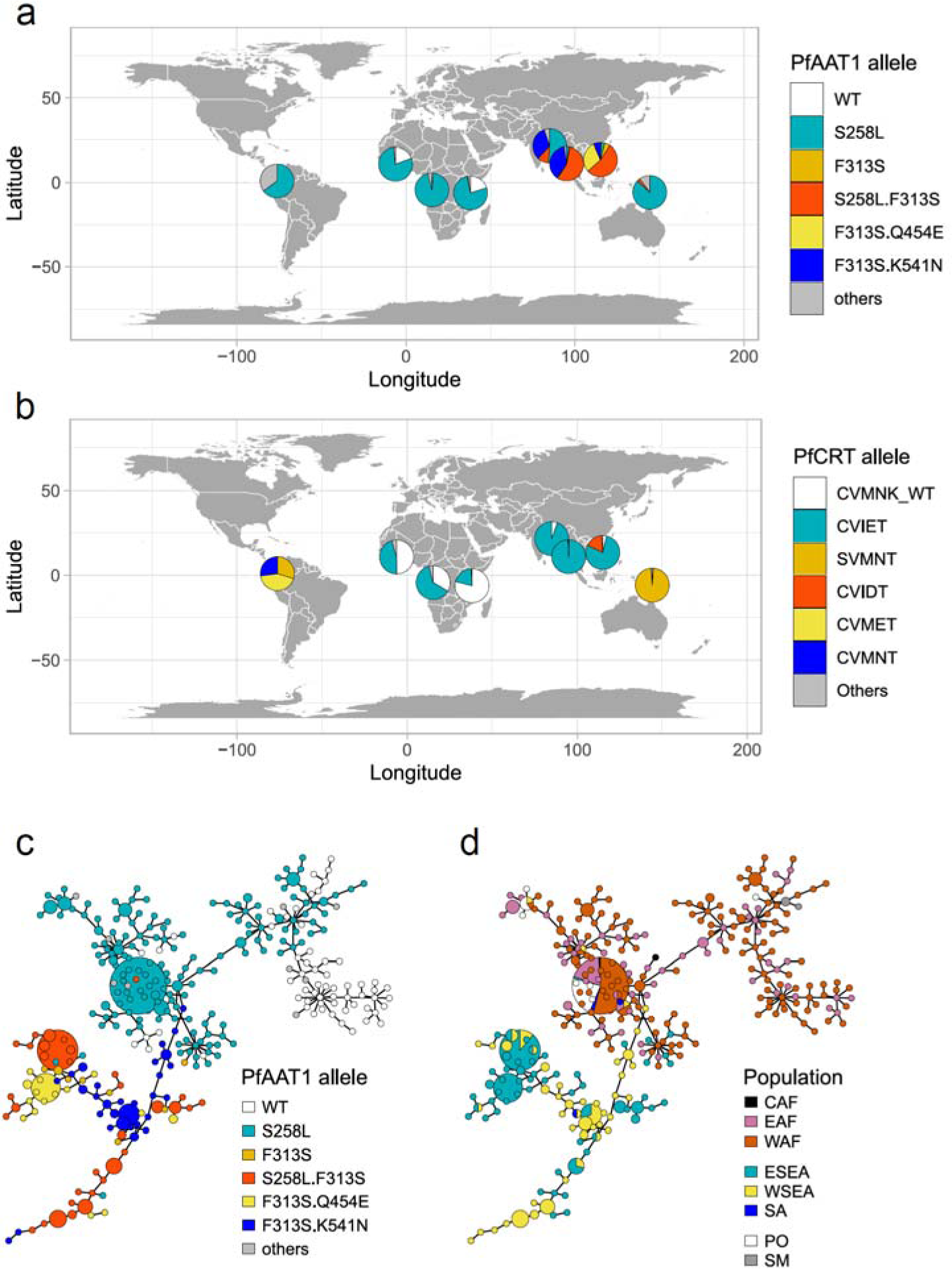
Distinctive trajectory of *pfaat1* evolution in Southeast Asia. (**a**) Global distribution of *pfaat1* alleles. (**b**) Comparable maps showing percentages of *pfcrt* haplotypes for amino acids 72-76. The colored segments show the major *pfcrt* haplotypes varying the K76<u>T</u> mutation. We used dataset from MalariaGEN release 6 for *pfaat1* and *pfcrt* allele frequency analysis. Data used for the figure is contained in Supplementary Table 2. Only samples with monoclonal infections (N = 4051) were included (1233 from West Africa [WAF], 415 from East Africa [EAF], 170 from Central Africa [CAF], 994 from east Southeast Asia [ESEA], 998 from west Southeast Asia [WSEA], 37 from South Asia [SA], 37 from South America [SM], and 167 from the Pacific Ocean region [PO]). (C and D) show minimum spanning networks of haplotypes colored by *pfaat1* allele (**c**) and geographical location (**d**), respectively. Networks were constructed from 50 kb genome regions centered by *pfaat1* (25 kb up- and down-stream. This spans the genome regions showing LD around *pfaat1* (Fig S5). 581 genomes with the highest sequence coverage were used to generate the network. The networks were generated based on 1847 SNPs (at least one mutant in the full dataset - MalariaGEN release 6). Circle size indicates number of samples represented (smallest, 1; largest, 87). Haplotypes from the same region (Asia or Africa) were clustered together, indicating independent origin of *pfaat1* alleles.

### Parasite genetic crosses using humanized mice identify a QTL containing pfaat1

*P. falciparum* genetic crosses can be achieved with human-liver chimeric mice, reviving and enhancing this powerful tool for malaria genetics ^23, 24^, after use of great apes for research was banned. We conducted three independent biological replicates of a cross between the CQ-sensitive West African parasite, 3D7, and a recently isolated CQ-resistant parasite from the Thailand-Myanmar border, NHP4026 (Supplementary Table 3). We then compared genome-wide allele frequencies in CQ-treated and control-treated progeny pools to identify quantitative trait loci (QTL). This bulk segregant analysis (BSA) ^25^ of progeny parasites robustly identified the chr. 7 locus containing *pfcrt* as expected, validating our approach (Fig. 3a and Supplementary Fig. 8 & 9). We were also intrigued to see a significant QTL on chr. 6 in each of the replicate crosses (Fig. 3, Supplementary Fig. 8-10). We prioritized genes within the 95% confidence interval of each QTL (Supplementary Table 4) by inspecting the SNPs and indels that differentiated the two parents (Supplementary Table 5). The chr. 6 QTL spanned from 1,013 kb to 1,283 kb (270 kb) and contained 60 genes. Of these, 54 are expressed in blood stages, and 27 have non-synonymous mutations that differentiate 3D7 from NHP4026. *pfaat1* was located at the peak of the chr. 6 QTL (Fig. 3c). NHP4026 carried two derived non-synonymous mutations in *pfaat1* (S258<u>L</u> and F313<u>S</u>) compared to 3D7, which carries the ancestral allele at each of these codon sites. We thus hypothesized that one or both of these *pfaat1* SNPs may be driving the chr. 6 QTL.

**Fig. 3.**
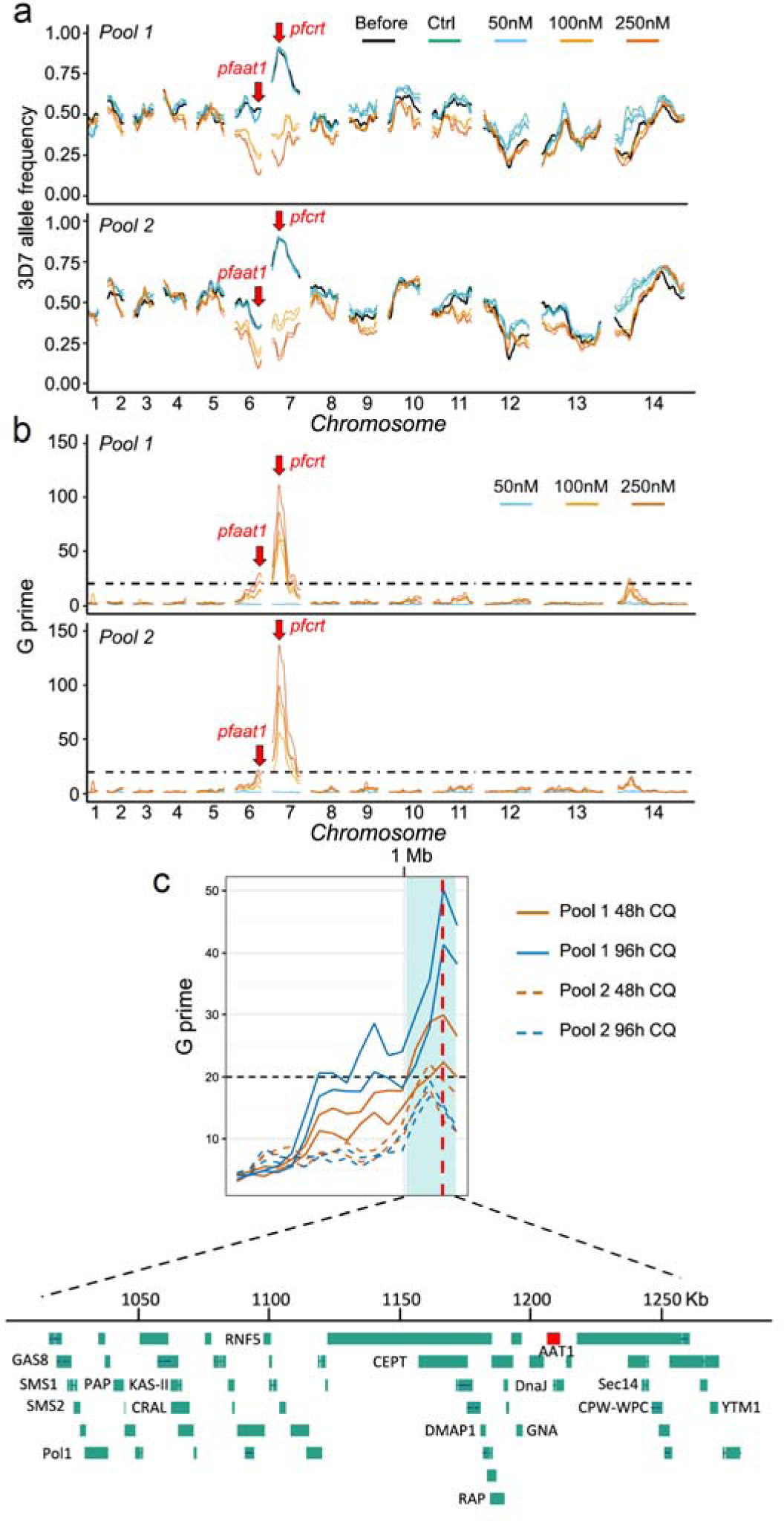
Genetic crosses and bulk segregant analysis reveal two QTL after CQ selection. (**a**) Allele frequency plots across the genome before and after CQ treatment. Lines with the same color indicate results from technical replicates. (**b**) QTLs identified using the G’ approach. Lines with the same color indicate results from technical replicates. Panels (**a**) and (**b**) include results from BSA with 48 hour CQ treatment with samples collected at day 4. See Figures S8 and S9 for the complete BSA from different collection time points and drug treatment duration under different CQ concentrations. (**c**) Fine mapping of the chr. 6 QTL. The 95% confidence intervals (CIs) were calculated from the 250 nM CQ treated samples, including data from different collection time points (day 4 for 48 hour CQ treatment and day 5 for 96 hour CQ treatment), pools (pool 1 and pool 2), and drug treatment duration (48 hour and 96 hour). Light cyan shadow shows boundaries of the merged CIs of all the QTLs. Each line indicates one QTL; black dashed line indicates threshold for QTL detection (*G prime* = 20). The vertical red dashed line indicates *pfaat1* location.

We isolated individual clones from the bulk 3D7 × NHP4026 F_1_ progeny to recover clones with all combinations of parental alleles at the chr. 6 and chr. 7 QTL loci. We cloned parasites both from a bulk progeny culture that was CQ-selected (96 hours at 250 nM CQ) and from a control culture. This generated 155 clonal progeny: 100 from the CQ-selected culture, 62 of which were genetically unique and 55 from the untreated control culture, of which 47 were unique (Fig. 4a). We compared allele frequencies between these two progeny populations (Fig. 4b), revealing significant differences at both chr. 6 and chr. 7 QTL regions, paralleling the BSA results. We observed a dramatic depletion of the NHP4026 CQ-resistant allele at the chr. 7 QTL in control treated cultures, consistent with strong selection against CQ resistant *pfcrt* alleles in the absence of CQ-selection. Conversely, all progeny isolated after CQ treatment harbored the NHP4026 CQ-resistant *pfcrt* allele. The inheritance of the *pfcrt* locus (chr. 7) and the *pfaat1* locus (chr. 6) was tightly linked in the isolated clones (Fig. 4c). To further examine whether the cross data were consistent with epistasis or co-selection, we examined a larger sample of recombinant clones isolated from five independent iterations of this genetic cross in the absence of CQ selection. This revealed significant under-representation of clones with genotype *pfcrt* 76<u>T</u> and *pfaat1* 258S/313F (WT) (Supplementary Table 6, X^2^ = 12.295, *p*-value = 0.0005). These results are consistent with the strong LD between these loci observed in nature (Supplementary Fig. 4) ^20^ and suggest a functional relationship between the two loci. A role for *pfaat1* S258<u>L</u>/F313<u>S</u> in compensating for the reduced fitness of parasites bearing *pfcrt* K76<u>T</u> is one likely explanation for the observed results.

**Fig. 4.**
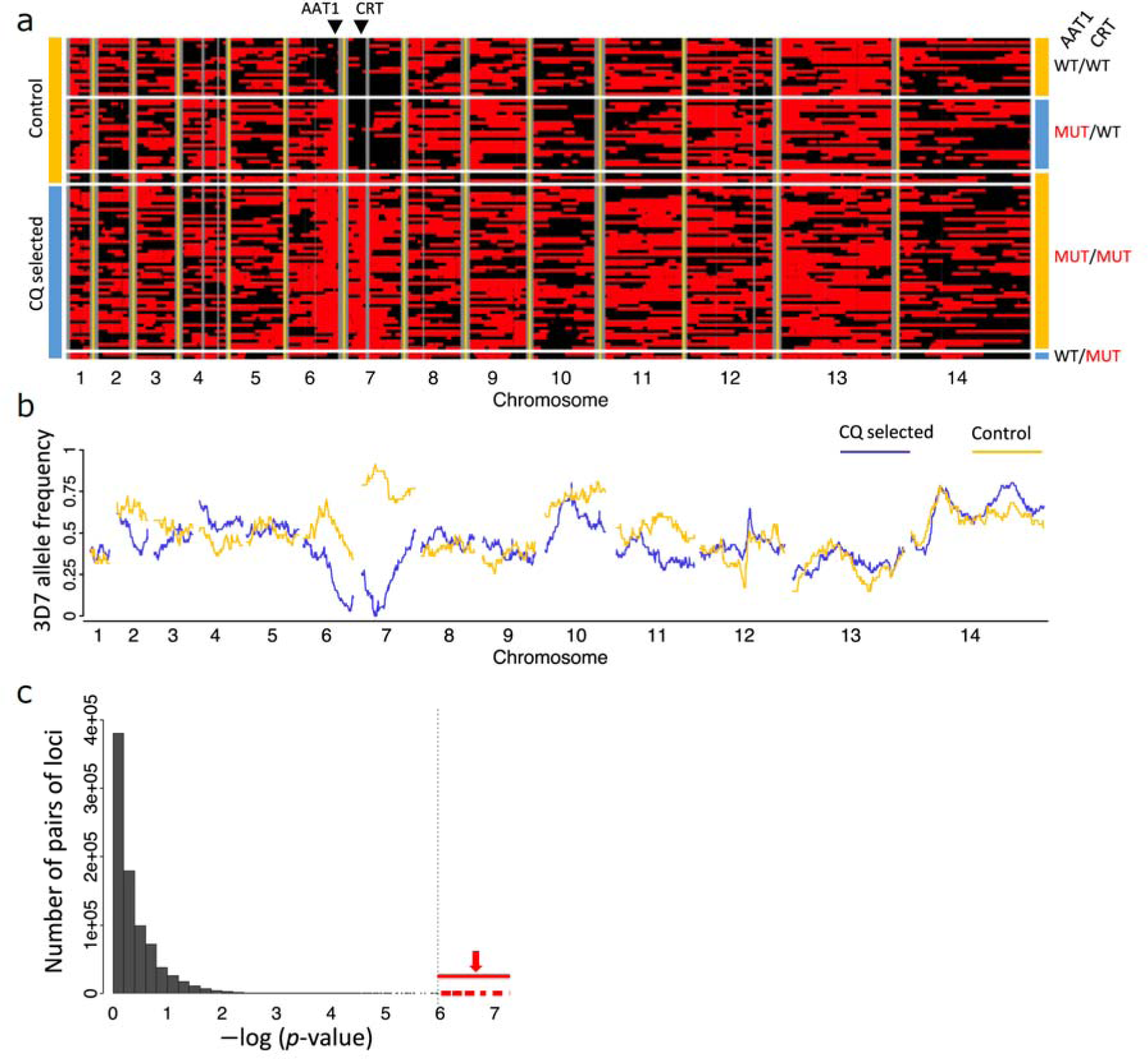
Analysis of cloned progeny reveals linkage and epistatic interactions between *pfcrt* and *pfaat1*. (**a**) Allelic inheritance of 109 unique recombinant progeny. Black and red blocks indicate alleles from 3D7 and NHP4026, separately. Vertical grey lines show non-core regions where no SNPs were genotyped. Clones isolated from recombinant progeny pools with or without CQ treatment are labeled on the left side of the panel. *pfaat1* and *pfcrt* alleles are labeled on the right side of the panel. WT indicates *pfaat1* and *pfcrt* alleles from 3D7 and MUT indicates alleles from NHP4026. The location of *pfaat1* and *pfcrt* are marked using black triangles on the top of the panel. (**b**) Genome-wide 3D7 allele frequency plot of unique progeny cloned from pools after 96-hours of CQ (250 nM) treatment (blue) or from control pools (gold). (**c**) Linkage between loci on different chromosomes measured by Fisher’s exact test. The dotted vertical line marks the Bonferroni corrected significance threshold, while points shown in red are comparisons between SNPs flanking *pfaat1* and *pfcrt*. Supplementary Table 6 shows non-random associations between genotypes in parasite clones recovered from untreated cultures.

We next measured *in vitro* CQ IC_50_ values for 18 parasites (a set of 16 progeny and both parents), carrying all combinations of the chr. 6 and chr. 7 QTL alleles (Supplementary Fig. 11, Supplementary Table 7). The NHP4026 parent was the most CQ-resistant parasite tested. All progeny that inherited NHP4026 *pfcrt* showed a CQ-resistant phenotype while all progeny that inherited 3D7 *pfcrt* were CQ-sensitive, consistent with previous reports. The effect *of pfcrt* alleles on parasite CQ resistance was significant based on a two-way ANOVA test (*p*=7.52 × 10^-11^). We did not see an impact of the *pfaat1* genotypes on IC_50_ in clones carrying *pfcrt* 76<u>T</u> (*p* = 0.06) or *pfcrt* 76K (*p* = 0.19). This analysis has limited power because only two progeny parasites were recovered with *pfaat1* 258S/313F (WT) in combination with *pfcrt* 76<u>T</u> (Fig.4a, Supplementary Fig. 11), but is consistent with the *pfAAT1* QTL being driven by parasite fitness in our genetic crosses. We therefore focused on gene manipulation of isogenic parasites for functional analysis.

### Functional validation of the role of pfaat1 in CQ resistance

We utilized CRISPR/Cas9 modification of the NHP4026 CQ-resistant parent to investigate the effects of mutations in *pfaat1* on CQ IC_50_ drug response and parasite fitness (Fig. 5). NHP4026 *pfaat1* carries the two most common SEA non-synonymous changes (S258<u>L</u> and F313<u>S</u>) (Fig. 2), relative to the sensitive 3D7 parent. We edited these positions back to the ancestral state both singly and in combination and confirmed the modifications in three clones isolated from independent CRISPR experiments for each allelic change (Fig. 5a). We then determined CQ IC_50_ and measured fitness using pairwise competition experiments for parental NHP4026^258<u>L</u>/313<u>S</u>^, the single mutations NHP4026^258<u>L</u>/313F^, NHP4026^258S/313<u>S</u>^ and the ancestral allele NHP4026^258S/313F^. This revealed a highly significant impact of the S258<u>L</u> mutation, which increased CQ IC_50_ 1.5-fold, and a more moderate but significant impact of F313<u>S</u> and the double mutation (S258<u>L</u>/F313<u>S</u>) (Fig. 5b, Supplementary Table 8). The observation that 258<u>L</u> shows an elevated IC_50_ only in combination with the ancestral 313F allele reveals an epistatic interaction between these amino acid variants (Fig 5B).

**Fig. 5.**
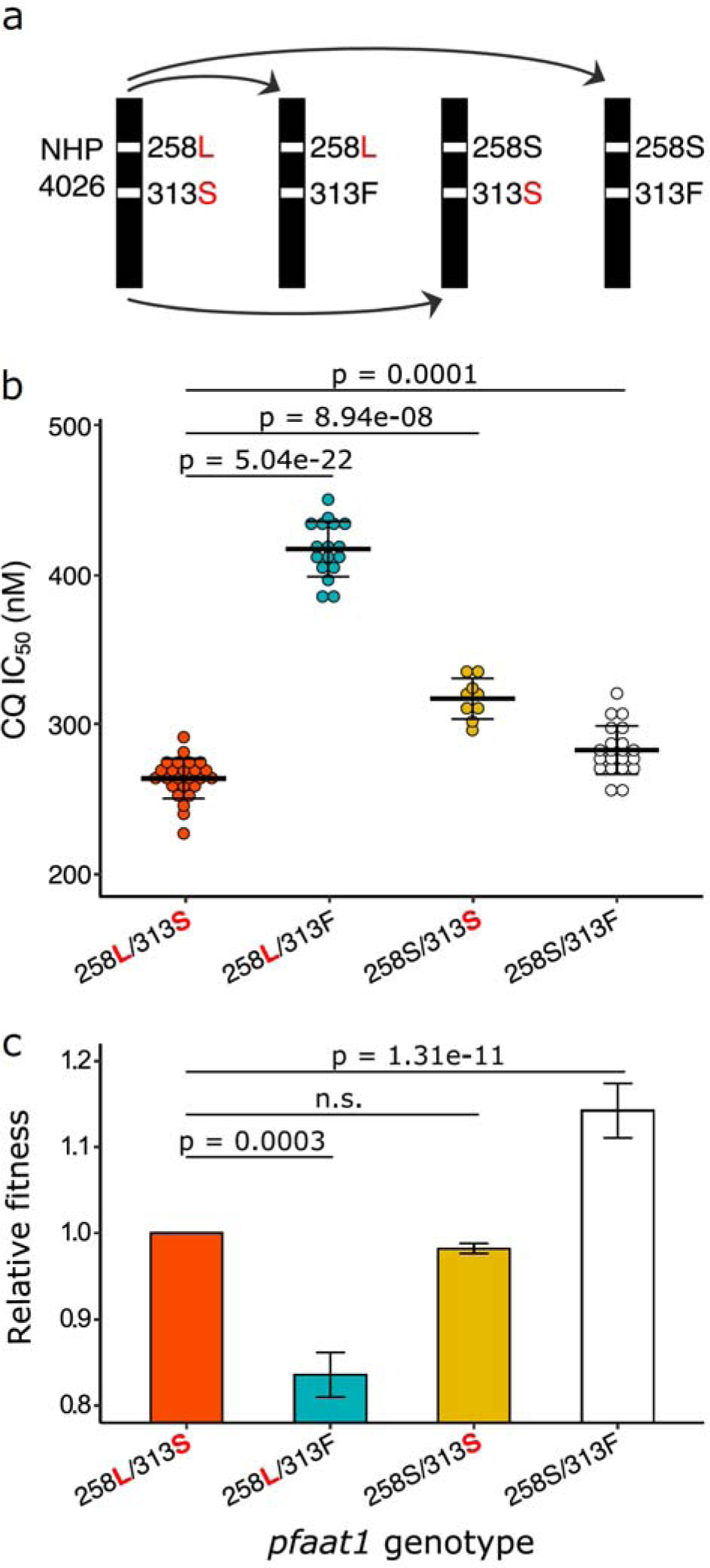
Allelic replacement impacts drug response and parasite fitness. (**a**) CRISPR/Cas9 gene editing. Starting with the NHP4026 parent, we generated all combinations of the SNP-states at *pfaat1*. (**b**) Drug response. Each dot indicates one replicate IC_50_ measurement: we used 2-4 independent CRISPR edited clones for each haplotype examined. Haplotypes are shown on the x-axis with derived amino acids shown in red. Bars show means (±1 s.e.), while significant differences between haplotypes are marked. (**c**) Fitness. The bars show relative fitness measured in replicated competition experiments conducted in the absence of CQ. See Fig S17 for allele frequency changes for each competition experiment. n.s., not significant.

We also examined the impact of the S258<u>L</u> and F313<u>S</u> substitutions on responses to other quinoline drugs. The results revealed significant impacts of PfAAT1 substitutions on quinine, amodiaquine and lumefantrine IC_50_ responses, and no impact on mefloquine IC_50_ (Supplementary Fig. 12). Notably, these IC_50_ value shifts were well below the threshold associated with clinical resistance. Consequently, although mutations in *pfaat1* can subtly impact susceptibly to a range of compounds, these results are consistent with CQ treatment being the primary selective force that drove the *pfaat1* S258<u>L</u> and F313<u>S</u> mutations along with those in *pfcrt*.

Mutations conferring drug resistance often carry fitness costs in the absence of drug treatment. We thus examined parasite fitness by conducting pairwise competition experiments with the parental NHP4026 parasite against the same mutant *pfaat1* parasites created above. This revealed significant differences in fitness (Fig. 5c). The 258<u>L</u>/313F allele that showed a selective sweep in The Gambia, was the least fit of all genotypes, the ancestral allele (258S/313F) carried by the 3D7 parent was the most fit, while the 258S/313<u>S</u> mutation, showed a similar fitness to the NHP4026 parent (258<u>L</u>/313<u>S</u>). These results also revealed strong epistatic interactions in fitness. While the 258<u>L</u>/313F allele that conferred high CQ IC_50_ (Fig. 5b), carried a heavy fitness penalty (Fig. 5c), fitness was partially restored by addition of the 313<u>S</u> mutation to generate the 258<u>L</u>/313<u>S</u> allele that predominates in SEA. Together these results show that the *pfaat1* S258<u>L</u> substitution underpins a 1.5-fold increase in CQ-resistance that likely drove its selective spread in The Gambia. However, S258<u>L</u> carries a high fitness cost that in SEA parasites was likely mitigated by addition of the compensatory substitution, F313<u>S</u>. Overall, these results demonstrate a large impact of *pfaat1* mutations on fitness of parasites carrying *pfcrt* K76<u>T</u> resistance alleles.

The editing experiments reveal that clones carrying the ancestral *pfaat1* allele in combination with *pfcrt* K76<u>T</u> show the highest fitness. In contrast, the close association of *pfaat1* S258<u>L</u>/F313<u>S</u> with *pfcrt* K76<u>T</u> in progeny from the genetic crosses revealed the opposite relationship. We speculate that these opposing results may reflect differing selection pressures in blood stage parasites in the case of CRISPR experiments, or in the mosquito and liver stages of the life cycle in the case of genetic crosses.

To further understand how *pfaat1* S258<u>L</u> impacts parasite phenotype, we used a yeast heterologous expression system. WT *pfaat1* is expressed in the yeast plasma membrane ^26^, where it increases quinine and CQ uptake conferring sensitivity to quinoline drugs, resulting in reduced growth. CQ uptake can be competitively inhibited by the aromatic amino acid tryptophan, suggesting a role for *pfaat1* in drug and amino acid transport ^26^. We therefore expressed *pfaat1* S258<u>L</u> in yeast, which restored yeast growth in the presence of high levels of CQ (Supplementary Fig. 13). Interestingly, expression of another amino acid variant (T162<u>E</u>), responsible for CQ-resistance in rodent malaria parasites (*Plasmodium chabaudi*) ^27^, also prevents accumulation of quinoline drugs within yeast cells and restores cell growth in the presence of CQ ^26^. Together, these new and published results suggest that yeast expression of *pfaat1* mutations impact resistance and fitness by altering the rates of amino acid and CQ transport.

We evaluated three-dimensional structural models based on the 3D7 PfAAT1 amino acid sequence using AlphaFold ^28^ and I-TASSER ^29^ (Supplementary Fig. 14). While CRT has ten membrane-spanning helices ^30^, AAT1 has eleven; this was corroborated using the sequence-based membrane topology prediction tool TOPCONS ^31^. The common AAT1 mutations S258<u>L</u>, F313<u>S</u> and Q454<u>E</u> are situated in membrane-spanning domains, while K541<u>L</u> is in a loop linking domains 9 and 10. The location of these high frequency non-synonymous changes in membrane spanning domains has strong parallels with CRT evolution ^30^ and is consistent with a functional role for these amino acids in transporter function.

## Discussion

Identification of *pfcrt* as the major determinant of CQ resistance was a breakthrough that transformed the malaria drug resistance research landscape, but the contribution of additional genetic factors in the evolution and maintenance of CQ resistance remained unclear ^32, 33^. By combining longitudinal population genomic analysis spanning the emergence of CQ resistance in The Gambia, analysis of bulk populations and progeny from controlled genetic crosses, and functional validation using both yeast and *P. falciparum*, we find clear evidence that a second locus, *pfaat1*, has played a central role in CQ resistance. This powerful combination of approaches allowed us to examine critical *pfaat1* variants that contribute to the architecture of CQ-resistance and interactions between *pfcrt* and *pfaat1*.

Our results provide compelling evidence that consolidates disparate observations from several systems suggesting a role for *pfaat1* in drug resistance evolution. In the rodent malaria parasite *P. chabaudi*, a mutation (T162<u>E</u>) in the orthologous amino acid transporter (*pcaat1*) was found to be a determinant of low level CQ resistance in laboratory-evolved resistance ^27^. In *P. falciparum* genome-wide association studies (GWAS), the S258<u>L</u> mutation of *pfaat1* was associated with CQ resistance in field isolates collected along the China-Myanmar border ^34^, while *pfcrt* K76<u>T</u> and *pfaat1* S258<u>L</u> show the strongest LD between physically unlinked chromosomes genome-wide ^20^. In addition, mutations in *pfaat1* have been linked to the *in vitro* evolution of resistance in *P. falciparum* to three different drug scaffolds ^35^. Previous work identified strong signatures of recent selection in parasites in Africa at regions surrounding *pfcrt*, *pfaat1*, and other drug resistance loci ^16, 17, 36^; similar signatures of selection are seen in Asia and South America ^18, 19^, while *pfaat1* was highlighted in a list of *P. falciparum* genes showing extreme geographical differentiation ^21^.

The different *pfaat1* haplotypes in Africa and Asia may be partly responsible for the contrasting evolution of CQ-resistance in these two continents. CQ-resistant parasites, carrying both *pfcrt* K76<u>T</u> and *pfaat1* S258<u>L</u> spread across Africa, but after removal of CQ as the first line drug, the prevalence of CQ resistant parasites declined in many countries ^37–39^. This is consistent with the low fitness of parasites carrying *pfcrt* K76<u>T</u> and *pfaat1* S258<u>L</u> in the absence of drug pressure, and intense competition within malaria parasite infection in Africa ^40^.

In contrast, *pfcrt* K76<u>T</u> has remained at or near fixation in many SEA countries ^21, 41^ (Fig. 2). On the Thailand-Myanmar border, CQ-resistance has remained at fixation since 1995, when CQ was removed as first line treatment of *P. falciparum* malaria ^41^. Our *pfaat1* mutagenesis results demonstrate that parasites bearing *pfaat1* 258<u>L</u>/313<u>S</u> show reduced IC_50_, but elevated fitness relative to *pfaat1* 258<u>L</u>/313F. The restoration of fitness by F313<u>S</u> may help to explain retention of CQ-resistant *pfcrt* K76<u>T</u> alleles in SEA. The alternative hypothesis – that high frequencies of F313<u>S</u> mutations are driven by widespread use of other quinoline partner drugs in SEA ^42^ is not supported, because we see only minor impacts of this substitution on response to lumefantrine, quinine, mefloquine and amodiaquine (Supplementary Fig. 12).

Mutations in *pfcrt* confer CQ-resistance by enabling efflux of CQ across the digestive vacuole membrane, away from its site of action ^8^. PfAAT1 is also located in the digestive vacuole membrane, where it likely acts as a bidirectional transporter of aromatic amino acids ^9, 43^. Given the structural similarity of quinoline drugs and aromatic amino acids, *pfaat1* mutations may modulate the ability of PfAAT1 to transport CQ and/or amino acids ^26, 43^. The *pfaat1* S258<u>L</u> mutation could potentiate resistance by either increasing efflux of CQ out of the digestive vacuole or reducing the rate of entry into the vacuole. Given that this *pfaat1* mutation blocks entry of quinoline drugs into yeast cells when heterologously expressed in the yeast cell membrane ^26^, we hypothesize that the *pfaat1* S258<u>L</u> mutation reduces CQ uptake into the food vacuole (Fig. 6). Our mutagenesis analyses show that the S258<u>L</u> allele has a high fitness cost, perhaps due to a decreased capacity for the amino acid transport from the vacuole. Interestingly, comparison of the *pfaat1* S258<u>L</u>/F313<u>S</u> haplotype segregating in our genetic cross with the WT *pfaat1* allele generated using gene editing, revealed only marginal increases in IC_50_ and limited reductions in fitness. This is consistent with the F313<u>S</u> mutation restoring the natural *pfaat1* function of transporting amino acids, thereby reducing osmotic stress and starvation, while also partially reducing levels of CQ-resistance (Fig. 6). That this haplotype has reached high frequency in SEA, may contribute to the maintenance of *pfcrt* K76<u>T</u> alleles long after the removal of CQ as a first line drug.

**Fig. 6.**
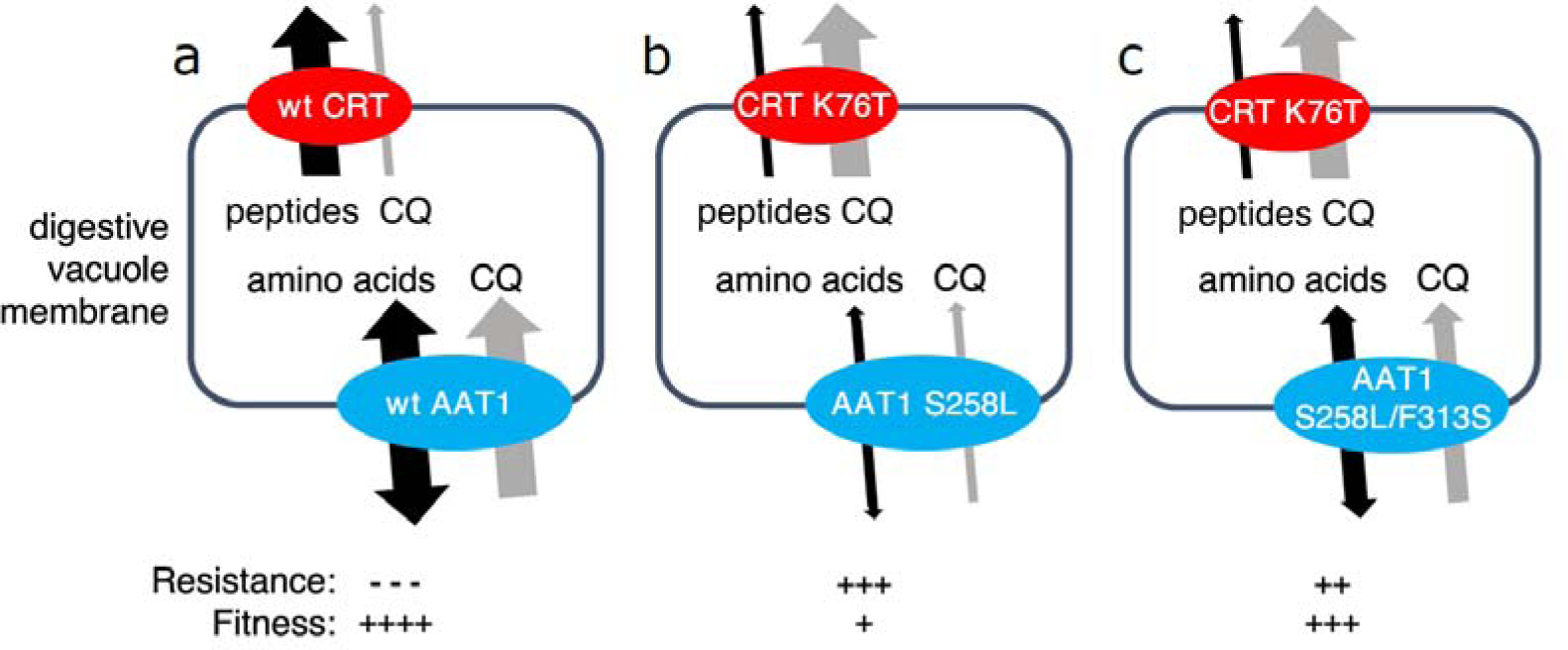
Model for involvement of *pfaat1* haplotypes in chloroquine resistance and fitness. PfCRT (red) and PfAAT1 (blue) are both situated in the digestive vacuole (DV) membrane. (**a**) Wildtype PfCRT and PfAAT1 transport peptides and aromatic amino acids respectively as well as chloroquine (CQ). (**b**) PfCRT K76<u>T</u> exports CQ from the DV away from its site of action, leading to elevated resistance but transports peptides inefficiently leading to a loss of fitness. PfAAT1 S258<u>L</u> reduces entry of CQ into the DV, leading to elevated resistance, but amino acid flux is affected leading to a loss of fitness. (**c**) The PfAAT1 S258<u>L</u>/F313<u>S</u> double mutation increases CQ influx in comparison to the S258<u>L</u> alone but the amino acid transport function is restored, leading to reduced IC_50_ and increased fitness in the absence of drug treatment.

Our results reveal hidden complexity in CQ-resistance evolution: drug treatment has driven global selective sweeps acting on mutations in an additional transporter (PfAAT1) located in the *P. falciparum* digestive vacuole membrane, which fine tune the balance between nutrient and drug transport, revealing evidence for epistasis and compensation, and impacting both drug resistance and fitness.

## Methods

### Ethics approval and consent to participate

The study was performed in strict accordance with the Guide for the Care and Use of Laboratory Animals of the National Institutes of Health (NIH), USA. The Seattle Children’s Research Institute (SCRI) has an Assurance from the Public Health Service (PHS) through the Office of Laboratory Animal Welfare (OLAW) for work approved by its Institutional Animal Care and Use Committee (IACUC). All of the work carried out in this study was specifically reviewed and approved by the SCRI IACUC.

### Project design

The project design is summarized in Supplementary Fig. 15. In brief, we use (i) population genomic analyses (ii) genetic crosses and quantitative genetics analysis followed by (iii) functional analyses to investigate the role of additional loci in CQ resistance.

### Gambia population analysis

#### P. falciparum genome sequences

*P. falciparum* infected blood samples collected from central (Farafenni) and coastal (Serrekunda) Gambia in 1984 and 2001 respectively, were processed for whole blood DNA and *P. falcip*arum genomes and deep sequenced at the Wellcome Trust Sanger Institute. Data from isolates collected from coastal Gambia in 2008 and 2014 had been published previously ^44, 45^ (see Supplementary Table 1 for details). Prior to sequencing, *P. falciparum* genomes were amplified from whole blood DNA of each sample from 1984 and 2001 using selective whole genome amplification (sWGA) and then subjected to paired-end short-read sequencing on the Illumina HiSeq platform ^46^. Short sequencing reads were mapped to the *P. falciparum* 3D7 reference genome using *bwa mem* (http://bio-bwa.sourceforge.net/). Mapping files (BAM) were sorted and deduplicated by Picard tools v2.0.1 (http://broadinstitute.github.io/picard/), and single-nucleotide polymorphism (SNP) and indel were called with GATK Haplotype Caller (https://software.broadinstitute.org/gatk/) following the best practices endorsed by the Pf3K project (https://www.malariagen.net/data/pf3K-5). Only SNP genotypes were used for this study. VCF files were generated by chromosome, and then were merged using *bcftools* (https://samtools.github.io/bcftools/bcftools.html) and filtered using *vcftools* (https://vcftools.sourceforge.net/). After filtration, only biallelic SNP variants with a VQSLOD score of at least 2, a map quality >30, and supported by no less than 5 reads per allelic variant were remained. SNPs with minor allele frequency <2% were further removed from our analysis. We also removed samples with > 10% genotypes missing. In the final dataset, there were in total of 16,385 biallelic SNP loci and 321 isolates (1984 [134], 1990 [13], 2001 [34], 2008 [75], and 2014 [65]) remained for further analysis. The complexity of infection (monogenomic or polygenomic) was estimated as the inbreeding coefficient *Fws* from the merged VCF file using R package *Biomix*. The short-read sequence data for all isolates and those from across Africa that have been previously published are available from the European Nucleotide Archive, and sample identifiers are given in Supplementary Table 1.

#### Allele frequencies and pairwise differentiation

For each sample with a complexity of infection greater than 1, the allele with most reads was retained for mixed-allele genotypes to create a virtual haploid genome variation dataset. Allele frequencies were calculated in plink and pairwise differences between temporal populations and genetic clusters were estimated by *Fst* using Weir and Cockerham’s method applied in the *hierfstat* package in R. The likelihood ratio test for allele frequency difference pFST was further calculated using *vcflib*. For a combined pFST *p*-value, the fisher method was performed in R *metaseq* package. The summary *p*-values were corrected for multiple testing using Benjamini-Hochberg (BH) method.

#### Genome scans for selection

We considered samples collected in the same year as a single population irrespective of the location of collection. We used the hapFLK approach to detect signatures of positive selection through haplotype differentiation following hierarchical clustering of Gambian temporal population groups compared to an outgroup from Tanzania, as described by Fariello et al., 2013 ^47^. *P*-values were computed for each SNP-specific value using the Python script provided with the hapFLK program, and values were corrected for multiple testing using the BH method. Secondly, we used pairwise relatedness based on identity by descent to derive an iR statistic for each SNP as implemented by the *IsoRelate* ^18^ package in R. Regions with overlapping iR and hapFLK -log10 *p*-values > 5 were considered as regions of interest.

### Population analysis on *pfaat1* and *pfcrt* evolution

#### Datasets

We included two datasets in this study: **1)** genotypes of 7,000 world-wide *P. falciparum* samples from MalariaGEN Pf community project (version 6.0) ^21^. This dataset includes samples from South America (SM), West Africa (WA), Central Africa (CA), East Africa (EA), South Asia (SA), the western part of Southeast Asia (WSEA), the eastern part of Southeast Asia (ESEA) and Pacific Oceania (PO). **2)** We also included 194 Thailand samples with whole genome sequencing data available from Cerqueira et al. ^15^, and merged them into the WSEA population. Duplicate sequences were removed according to the sample’s original ID (Hypercode). Only samples with single parasite infections (within-host diversity *F_WS_* > 0.90) and >50% of SNP loci genotyped were included for further analysis. A total of 4051 samples remained after filtration (Supplementary Table 2). Non-biallelic SNPs and heterozygous variant calls were further removed from the dataset. We then extracted genotype data at *pfaat1* and *pfcrt* gene regions and calculated the allele frequencies (Fig 2A).

#### Pfaat1 haplotypes and evolutionary relationships

To minimize the effect from recombination, we extracted 1847 SNPs distributed within 25kb upstream and 25kb downstream of the *pfaat1* gene. For clarity, only samples with all 1847 SNPs genotyped (581/4051) were used for evolutionary analysis. To visualize the population structure, we calculated the genetic distance between every two samples and generated a minimum spanning network (MSN, Fig 2B and Supplementary Fig. 4), using R package *poppr*. We compared genome sequences (PlasmoDB, version 46) between *P. falciparum* and *Plasmodium reichenowi* and extracted genotypes at 1803/1847 common loci. We then built a UPGMA tree rooted by *P. reichenowi* using the 581 haplotypes and 1803 SNPs (Supplementary Fig. 7), using the R packages *ape* and *phangorn* under default parameters. MSN network and UPGMA tree were plotted with ggplot2.

### Genetic cross and bulk segregant analysis

#### Genetic cross preparation

We generated genetic crosses between parasite 3D7 and NHP4026 ^48^, using FRG NOD huHep mice with human chimeric livers and *A. stephensi* mosquitoes as described previously ^23–25, 49, 50^. 3D7 has been maintained in the lab for decades and is CQ sensitive; while NHP4026 was recently cloned from a patient visiting the Shoklo Malaria Research Unit (SMRU) clinic on the Thailand-Myanmar border in 2007 and is CQ resistant (Supplementary Table 3). For the genetic cross 3D7×NHP4026, we generated three recombinant pools using independent cages of infected mosquitoes: these are independent pools of recombinants ^48^. The estimated number of recombinant genotypes in each pool was ∼2800 ^48^. We used two of the pools (pool 1 and pool 2) that were maintained in culture media containing AlbuMAX for this study.

#### Drug treatment and sample collection

For each recombinant pool, the parasite culture was expanded under standard culture conditions as described in Brenneman et al. ^25^. Briefly, cultures were maintained in complete media (CM) at 5% hematocrit in O^+^ red blood cells (RBC) (Biochemed Services, Winchester, VA) at 37°C between a pH of 7.0-7.5 in an atmosphere of 5% CO_2_, 5% O_2_ and 90% N_2_. Media changes were performed every 48 hours and cultures were expanded to keep the parasitemia at ∼1%. Once expanded sufficiently, each recombinant pool was divided into sixteen 0.5 ml aliquots while diluting to 1% parasitemia. The aliquots were then maintained in 48-well plates and treated with CQ (Supplementary Fig. 16). In total, we had 32 cultures: 2 pools × 4 CQ concentrations (0 [control], 50, 100 or 250 nM) × 2 drug duration time (48 hour or 96 hour) × 2 technical replicates. We define the day when drug was applied as day 0. After two days (48 hour) of drug treatment, the infected RBCs were washed with Phosphate-buffered saline (PBS) solution twice to remove residual drug. For the plate assigned for 48 hour CQ treatment (48-well plate 1), cultures were then maintained in complete media (CM); and samples were collected at day 0, 4 and 7. For the plate assigned for 96 hour CQ treatment (48-well plate 2), fresh CQ was added back to the culture media and treated for another 48 hour; and after a total of 96 hour CQ treatment, drug was removed and samples were collected at day 0, 5 and 10. CQ was dissolved in H_2_O and diluted in incomplete media (ICM) (Gibco, Life Technologies). Culture medium was changed every 48 hours. Parasitemia was monitored using 20% Giemsa-stained slides and cultures were diluted to 1% parasitemia if the parasitemia was higher than 1%. Approximately 15 _μ_l packed RBCs was collected per sample.

#### Library preparation and sequencing

We prepared Illumina libraries and sequenced both parents and the 96 segregant pools collected above. We extracted and purified genomic DNA using the Qiagen DNA mini kit and quantified the amount of DNA with Quant-iT™ PicoGreen® Assay (Invitrogen). For samples with less than 50 ng DNA obtained, we performed whole genome amplification (WGA) following Nair et al. ^51^. WGA products were cleaned with KAPA Pure Beads (Roche Molecular Systems, Inc.) at a 1:1 ratio. We prepared sequencing libraries using 50-100 ng DNA or WGA product using KAPA HyperPlus Kit following the instruction with 3-cycles of PCR. All libraries were sequenced at 150bp pair-end using Illumina Novaseq S4 or Hiseq X sequencers, to get >100× genome coverage per sample.

#### Mapping and genotyping

We mapped the sequencing reads against the 3D7 reference genome (PlasmoDB version 46) using BWA mem (http://bio-bwa.sourceforge.net/), and deduplicated and trans-formatted the alignment files using picard tools v2.0.1 (http://broadinstitute.github.io/picard/). We recalibrated the base quality score based on a set of verified known variants ^52^ using *BaseRecalibrator*, and called variants through *HaplotypeCaller*. Both functions were from Genome Analysis Toolkit GATK v3.7 (https://software.broadinstitute.org/gatk/). Only variants located in the core genome regions (defined in ^52^) were called and used for further analysis.

#### Genotype of parents

We merged calls from the two parents using *GenotypeGVCFs* in GATK, and applied *standard* filtration to the raw variant dataset as described in ^53^. We recalibrated the variant quality scores (VQSR) and removed loci with VQSR < 1. The final variants in VCF format were annotated using snpEff v4.3 (https://pcingola.github.io/SnpEff/) with 3D7 (PlasmoDB, release46) as the reference. After filtration and annotation, we selected SNP loci that are distinct in the two parents and used those SNPs for further bulk segregant analysis.

#### Bulk segregant analysis

We used statistical methods described previously in ^25, 48, 50^ for bulk segregant analysis. The variant calls from segregant progeny pools were merged together. Additionally, SNP loci with coverage < 30× were removed. We counted reads with genotypes of each parent and calculated allele frequencies. Allele frequencies of 3D7 were plotted across the genome, and outliers were removed following Hampel’s rule ^54^ with a window size of 100 loci. We performed the bulk segregant analysis using the R package *QTLseqr* ^55^. Extreme-QTLs were defined as regions with G prime > 20 ^56^. Once a QTL was detected, we calculated an approximate 95% confidence interval using Li’s method ^57^ to localize causative genes.

### Progeny cloning and phenotyping

#### Progeny cloning

Individual progeny were cloned via limiting dilution at 0.3 cells per well from bulk cultures on day 10 after 96 hours of control/250nM CQ treatment. Individual wells with parasites were determined by qPCR (as described in Button-Simons et al.^49^) and expanded to larger cultures under standard culture conditions to obtain enough material for both cryopreservation and genome sequencing.

#### Sequencing and genotyping

Cloned progeny were sequenced and genotyped as described above in the <u>Genetic cross and bulk segregant analysis</u> section, with these modifications: 1) the cloned progeny were sequenced at 25× genome coverage; 2) SNP calls were filtered out if the coverage was < 3 reads per sample.

#### Cloned progeny analysis

Unique recombinant progeny were identified from all cloned progeny using a pipeline described in Button-Simons et al.^49^. Non-clonal progeny were identified based on the number and distribution of heterozygous SNP calls. Selfed progeny were identified as having greater than 90% sequence similarity to either parent. Unique recombinant progeny that were sampled multiple times during cloning were identified as clusters of individual clonal progeny with greater than 90% sequence similarity. We plotted frequencies of 3D7 alleles across the genome in progeny populations with and without CQ treatment. Heatmaps were generated to visualize inheritance patterns in individual unique recombinant progeny (Fig 4A). We selected sixteen unique recombinant progeny with different allele combination at chromosome 6 and chromosome 7 QTL regions for further CQ IC_50_ measurement (Fig S11).

#### Genome-wide linkage analysis on pfaat1 in cloned progeny

Fisher’s exact test was used to test for linkage between all inter-chromosomal pairs of loci across the set of 109 unique recombinant progeny. The distribution of the -log of the resulting *p*-values was plotted in Fig 4C and the significance cut-off was calculated based on a Bonferroni correction for the number of loci.

#### IC_50_ measurement for cloned progeny

Cryopreserved stocks of 3D7, NHP4026, 3D7×NHP4026 progeny were thawed and grown in CM under standard culture conditions as described above. Cultures were kept below 3% parasitemia with media changes every 48 hours. Parents and progeny IC_50_ was assessed via a standard 72 hour SYBR Green 1 fluorescence assay^58^. Cultures were assessed daily for parasitemia and stage. Cultures that were at least 70% ring were loaded into CQ dose-response assays of a series of 2 fold drug dilutions across 10 wells at 0.15% parasitemia. Drug stocks (1mg/ml) for CQ were prepared in H_2_O as single-use aliquots and stored at -20°C until use. Drug dilutions were prepared in incomplete media. Biological replicates were conducted with at least two cycles of culturing between load dates. IC_50_ values were calculated in GraphPad Prism 8 using a 4 parameter curve from two technical replicates loaded per plate.

### CRISPR/Cas9 editing at *pfaat1* and parasite phenotyping

#### CRISPR/Cas9 editing

We designed plasmids for CRISPR/Cas9 editing using the approach described in ^59^. The guide RNA (*GAAATTAAATACATAAAAGA*) was designed to target *pfaat1* in NHP4026. Edits (258<u>L</u>/313F, 258S/313<u>S</u> and 258S/313F, Fig. 5A) were introduced to NHP4026 through homology arm sequence with target and shield mutations. The parasites were transfected at ring stages with 100 µg plasmid DNA, and successful transfectants were selected by treatment with 24 nM WR99210 (gifted by Jacobus Pharmaceuticals, Princeton, NJ) for 6 days. The parasites were recovered after ∼3 weeks. To determine if recovered parasites contained the expected mutations, we amplified the target region (forward primer, AGTACGGTACTTTTTATATGTACAGCT; reverse primer, TGCATTTGGTTGTTGAGAGAAGG) and confirmed the mutation with Sanger sequencing. We cloned parasites from successful transfection experiments: independent edited parasites (from different transfection experiments) were recovered for each *pfaat1* genotype. Edited parasites were then genome sequenced to identify off-target edits elsewhere in the genome. We were not able to find any SNP or indel changes between the original NHP4026 and any CRISPR edited parasites other than the target and shield mutations.

#### IC_50_ measurement for CRISPR/Cas9 edited parasites

Parasite IC_50s_ for CQ, AMD, LUM, MQ and QN were measured for 3-5 clones per CRISPR/Cas9 modified line and for NHP4026 across multiple load dates as described above for cloned progeny, except that each plate included two NHP4026 technical replicates as controls. This replication of genotype within each load date allowed for detection of batch effects due to load date.

#### Batch correction for IC_50_ data

Analysis of Variance (ANOVA) was used to account for batch effects and to test for differences in IC_50_ between all genotype groups and for each contrast between each CRISPR/Cas9 modified line and NHP4026 for each drug tested ^60^. A linear model with load date (batch) and genotype as explanatory variables was utilized to generate batch corrected IC_50_ values for visualization of the impact of CRISPR/Cas9 modifications (Fig 5B and Supplementary Fig. 12).

#### Measurement of parasite fitness using competitive growth assays

Parasites were synchronized to late stage schizonts using a density gradient ^61^. The top layer of late stage schizonts was removed and washed twice with RPMI. Synchronized cultures were suspended in 5 ml of CM at 5% hematocrit and allowed to reinvade overnight with gentle shaking. Parasitemia and parasite stage were quantified using flow cytometry. Briefly, 80 _μ_l of culture and an RBC control were stained with SYBR Green I and SYTO 61 and measured on a guava easyCyte HT (Luminex Co.). 50,000 events were recorded to determine relative parasitemia and stage. When 80% of parasites were in the ring stage, the head-to-head competition experiments were set-up as previously described ^62^. Competition assays were set up between CRISPR/Cas9 edited parasites and NHP4026 in a 1:1 ratio at a parasitemia of 1% in a 96-well plate (200 _μ_l per well) and maintained for 30 days. Each of the assays contained three biological replicates (three independent clones from different CRISPR/Cas9 editing experiments) and two technical replicates (two wells of culture). Every two days, the parasitemia was assessed by microscopy using Giemsa-stained slides, samples were taken and stored at -80°C and the cultures were diluted to 1% parasitemia with fresh RBCs and media. The proportion of parasite in each competition (Supplementary Fig. 17) was measured using a rhAmp SNP Assay (IDT, Integrated DNA Technologies, Inc.) with primers targeting the CRISPR/Cas9 edited region in *pfaat1*.

#### Selection coefficient

We measured selection coefficient (*s*) by fitting a linear model between the natural log of the allele ratio (freq [allele edited parasite]/freq [NHP4026]) and time (measured in 48 hour parasite asexual cycles). The slope of the linear model provides a measure of the driving *s* of each mutation ^63^. To compare relative fitness of parasites carrying different *pfaat1* alleles, we normalized the fitness of NHP4026 to 1 and used slope + 1 to quantify the fitness of CRISPR/Cas9 edited parasites (Fig 5C).

### Overexpression of PfAAT1 in yeast

To generate pfAAT1 expressing yeast, plasmid carrying the *pfaat1* coding sequence was transformed into yeast *Saccharomyces cerevisiae* (BY4743) as described in ^26^. The doubling time (h) was measured for strains carry empty vector, wild-type pfAAT1, or S258<u>L</u> mutant pfAAT1. We measured doubling time under two culture conditions: control or with 5 mM CQ. Three independent experiments were performed for each assay.

### PfAAT1 protein structure analysis

Three-dimensional homology models for PfAAT1 were predicted using AlphaFold ^28, 64^ and I-TASSER ^29, 65, 66^ and analyzed with PyMol software (v2.3.0; Schrödinger, LLC). At the primary sequence level, we used TOPCONS ^31^ to predict transmembrane helix topology for comparison. We plotted a cartoon version of the protein transmembrane topology based on the computationally predicted structures and membrane topology (Supplementary Fig. 14). Models were truncated to exclude amino-terminal residues 1-166, likely positioned outside of the membrane, because AlphaFold assigns low confidence to this N-terminal stretch. Furthermore, mutations of interest map only to transmembrane helices according to both 3-D models and TOPCONS. I-TASSER generated models with topology similar to AlphaFold with the highest variations in AlphaFold low-confidence regions 1-166 and 475-516. The top five I-TASSER models superimpose on the AlphaFold model with a root-mean-square-deviation (RMSD) range of 2.4-2.8 Å over 303-327 of 440 aligned residues using the PDBeFold Server (http://www.ebi.ac.uk/msd-srv/ssm). The four common SNPs (S258<u>L</u>, F313<u>S</u>, Q454<u>E</u> and K541<u>L</u>) overlay closely between the homology models. We evaluated the impact of different mutations on protein stability using the mutagenesis function in PyMol.

### Data Availability

All raw sequencing data have been submitted to the NCBI Sequence Read Archive (SRA, https://www.ncbi.nlm.nih.gov/sra) or European Nucleotide Archive (ENA) with accession numbers available in supplementary Table 1. All other data are available in the main text or supplementary materials. The code used in analysis and data analyzed are available at GitHub through the following links: https://github.com/emilyli0325/CQ.AAT1.git (XL), https://github.com/MPB-mrcg?tab=repositories(AAN), and https://github.com/kbuttons/CQ.AAT1.progeny.git (KBS).

## Acknowledgments

We thank past and present staff and visiting researchers at the MRC Unit in The Gambia involved in studies from which samples were archived, including those under the earlier directorships of Prof Brian Greenwood and Prof Tumani Corrah. Work at Texas Biomedical Research Institute was conducted in facilities constructed with support from Research Facilities Improvement Program grant C06 RR013556. SMRU is part of the Mahidol Oxford University Research Unit supported by the Wellcome Trust of Great Britain. We thank the staff of the Wellcome Trust Sanger Institute Sample Logistics, Sequencing and Informatics facilities for their contributions. The Structural Biology Core at The University of Texas Health Science Center at San Antonio is a part of Institutional Research Cores supported by the Office of the Vice President for Research and the Mays Cancer Center Drug Discovery and Structural Biology Shared Resource (National Institutes of Health grant P30 CA054174).

## Author Contributions

AAN, KBS, XL, SK, and KVB contributed equally to this work.

Conceptualization: MTF, TA, IC, AV, AAN, DC

Methodology: AV, SK, SVA, ABT

Investigation: AAN, UDA, DK, DC, RA, RDP, KBS, KVB, LC, HD, JKL, AR, ED, SK, DS, ST, MF, ABT, TA

Analysis: XL, KBS, AAN

Visualization: XL, KBS, AAN

Funding acquisition : MTF, TA, DC, AAN, DK, SVA, FN

Project administration: MF, DK, AAN, DC, SVA

Supervision: MF, AV, TA, IC, SHIK, SVA, DK, DC

Writing – original draft: KVB, TA, KBS, XL

Writing – review & editing: MTF, AV, IC, SK, AAN, DC, DK, UDA, RDP, SHIK, FN, ABT, TA

## Competing Interests

Authors declare that they have no competing interests.

## Additional Information

Supplementary Information is available for this paper.

Correspondence and requests for materials should be addressed to Timothy J. C. Anderson.

Reprints and permission information is available at www.nature.com/reprints.

## Funding

National Institutes of Health grant P01 12072248 (MTF)

National Institutes of Health grant R37 AI048071 (TA)

MRC Career Development Fellowship MC_EX_MR/K02440X/1 (AAN)

MRC malaria program grant MC_EX_MR/J002364/1 (UDA)

European Research Council grant AdG-2011-294428 (DC and AAN)

MRC grant MR/S009760/1 (DC)

Wellcome Trust (206194, 204911) and the Bill & Melinda Gates Foundation (OPP1204628, INV-001927) (DK)

Biotechnology and Biological Sciences Research Council grant BB/M022161/1 (SVA)

Wellcome Trust [Grant number 220221] (FN); For the purpose of open access, the author has applied a CC BY public copyright license to any Author Accepted Manuscript version arising from this submission.

## Supplementary Figures

**Fig. S1.**
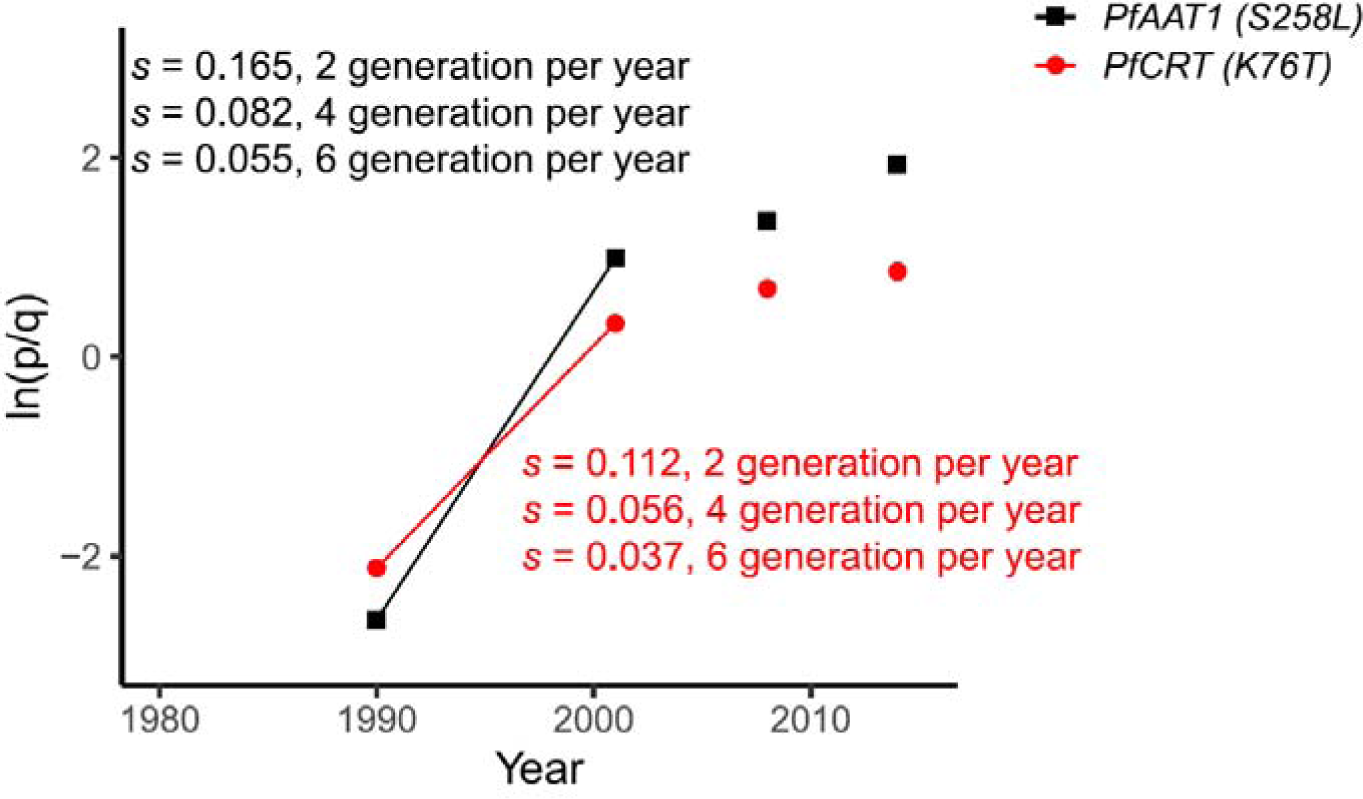
Estimation of selection coefficient (*s*) for *pfaat1 (S258<u>L</u>)* and *pfcrt (K76<u>T</u>)* alleles. *p* is the frequency of mutant alleles (*pfaat1* S258<u>L</u> or *pfcrt* K76<u>T</u>) as indicated in Fig. 1A, and q (=1-p) is the inferred frequency of wild-type (3D7) alleles. The x-axis indicates parasite generations (labeled with sample collection year). We estimated selection coefficients (*s*) based on allele frequency from year 1990 and 2001, as CQ monotherapy was stopped in Gambia in 2004. *s* indicates the changes in relative growth per parasite generation (i.e. the duration of the complete lifecycle in both mosquito and human host). The calculation was based on estimates of 2, 4, or 6 generations per year.

**Fig. S2.**
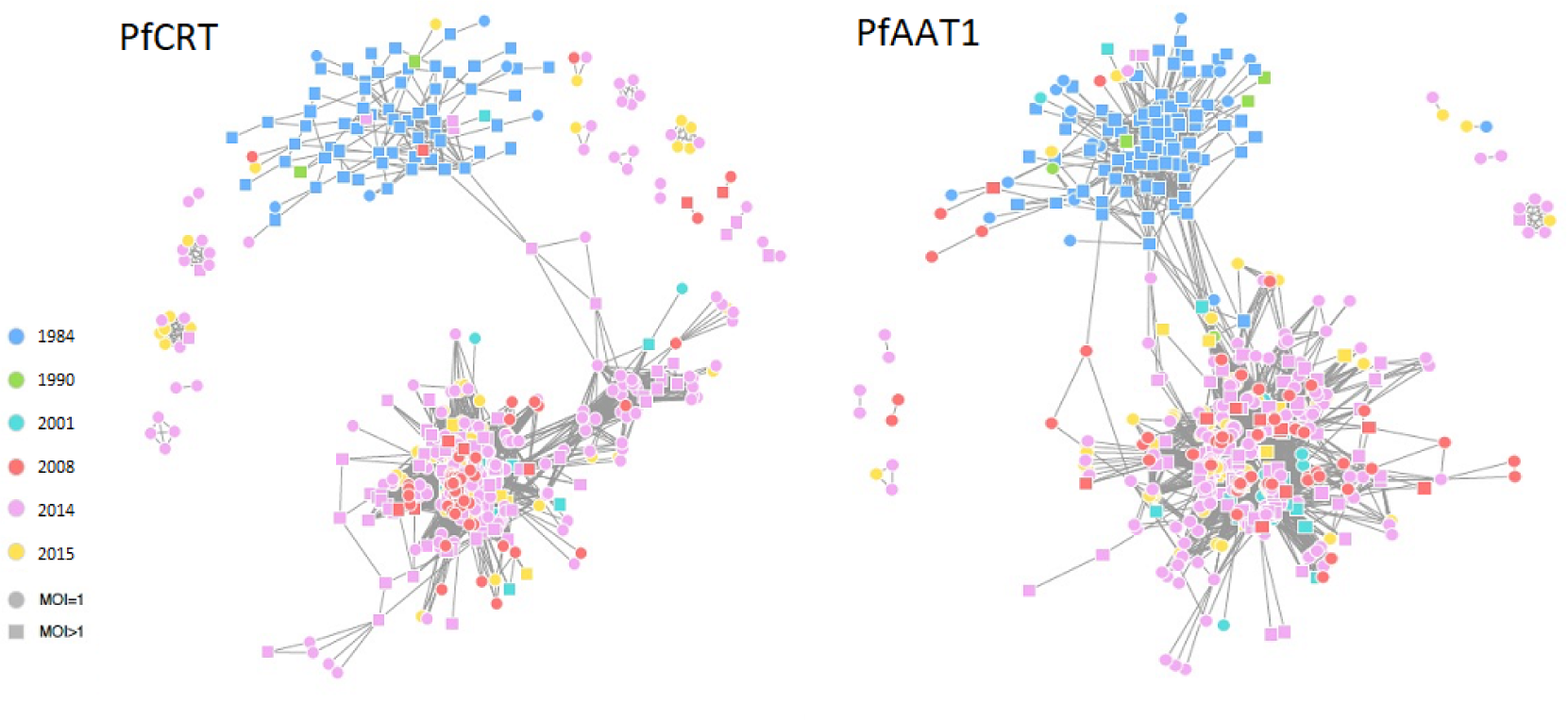
Haplotype structure at the *pfcrt* (left panel) and *pfaat1* (right panel) regions. Each point depicts an isolate with point colors representing the years from which they were sampled. Square points represent complex infections and circles represent monoclonals. MOI, multiplicity of infection.

**Fig. S3.**
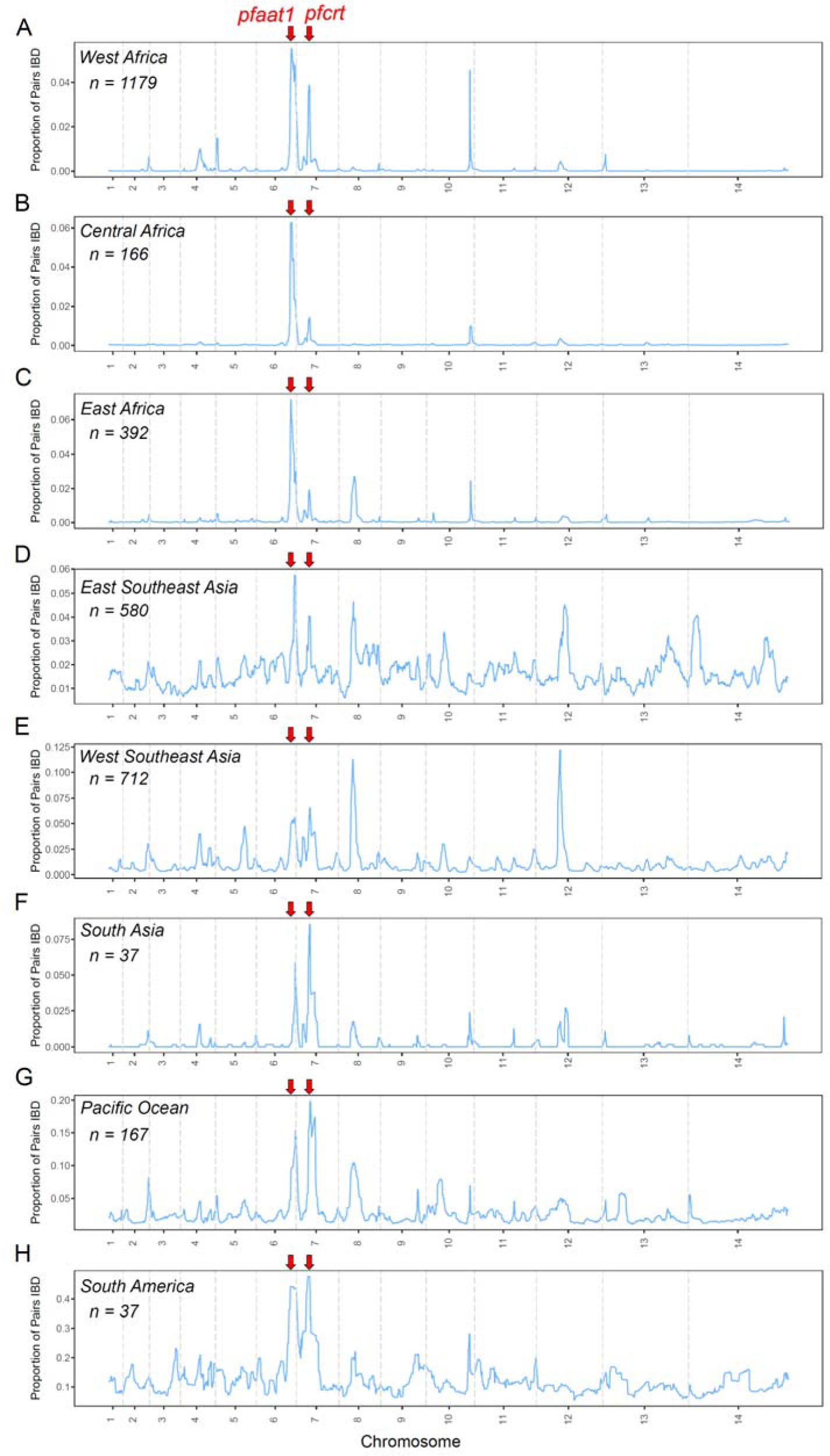
The proportion of pairs identical by descent (IBD) within populations from global locations. For samples with >90% of the genomes are IBD, only one representative sample with the highest genotype rate was selected and used for IBD analysis. Sample numbers are showed in each panel. Chromosome boundaries are indicated with grey dashed vertical lines. The location of *pfaat1* and *pfcrt* are indicated with red arrows on top of each panel. See also analysis by Amambua-Ngwa et al ^17^, Hendon et al ^18^, and Carrasquilla et al ^19^.

**Fig. S4.**
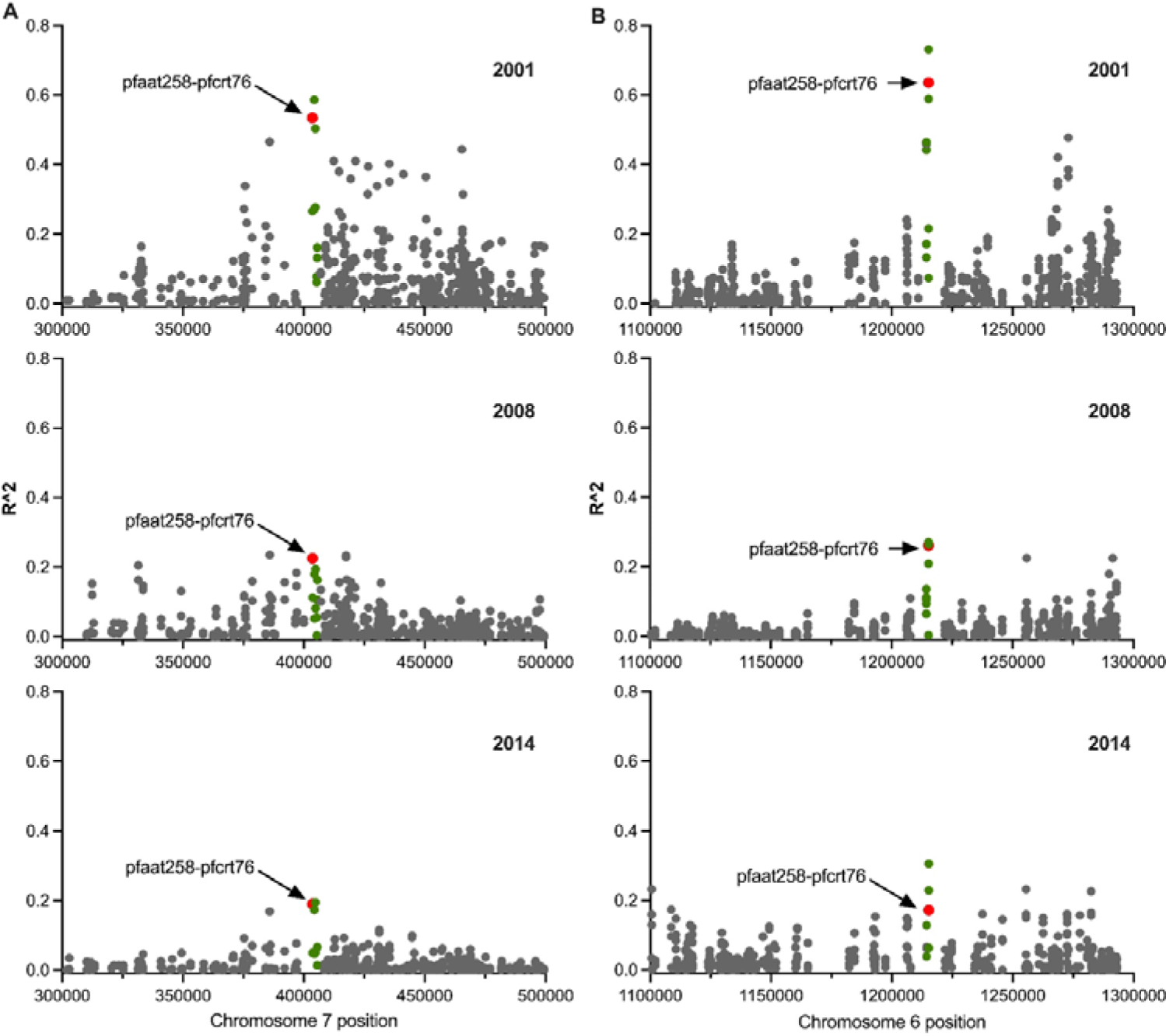
Interchromosomal linkage disequilibrium (LD) analysis. A. *R^2^* between *pfaat1* (SNP pfaat258 located at Pf3D7_06_v3:1215233) and a 200 kb region around the *pfcrt* gene on chr. 7. B. *R^2^* between *pfcrt* (SNP pfcrt76 located at Pf3D7_07_v3:403625) and a 200kb region around the *pfaat1* gene on chr. 6. *R^2^* values against SNPs within gene *pfcrt* (in panel A) or *pfaat1* (in panel B) are shown as green or red points, while the red points are LDs between SNPs pfaat258 and pfcrt76. For both panel A and B, from top to bottom, rows represent LD analysis for populations from year 2001, 2008, and 2014, respectively. Samples from 1984 are not shown, because neither *pfaat1* S258<u>L</u> nor *pfcrt* K76<u>T</u> were present. 1990 was omitted because only 13 samples were available. The highest LDs were observed between gene *pfaat1* and *pfcrt*.

**Fig. S5.**
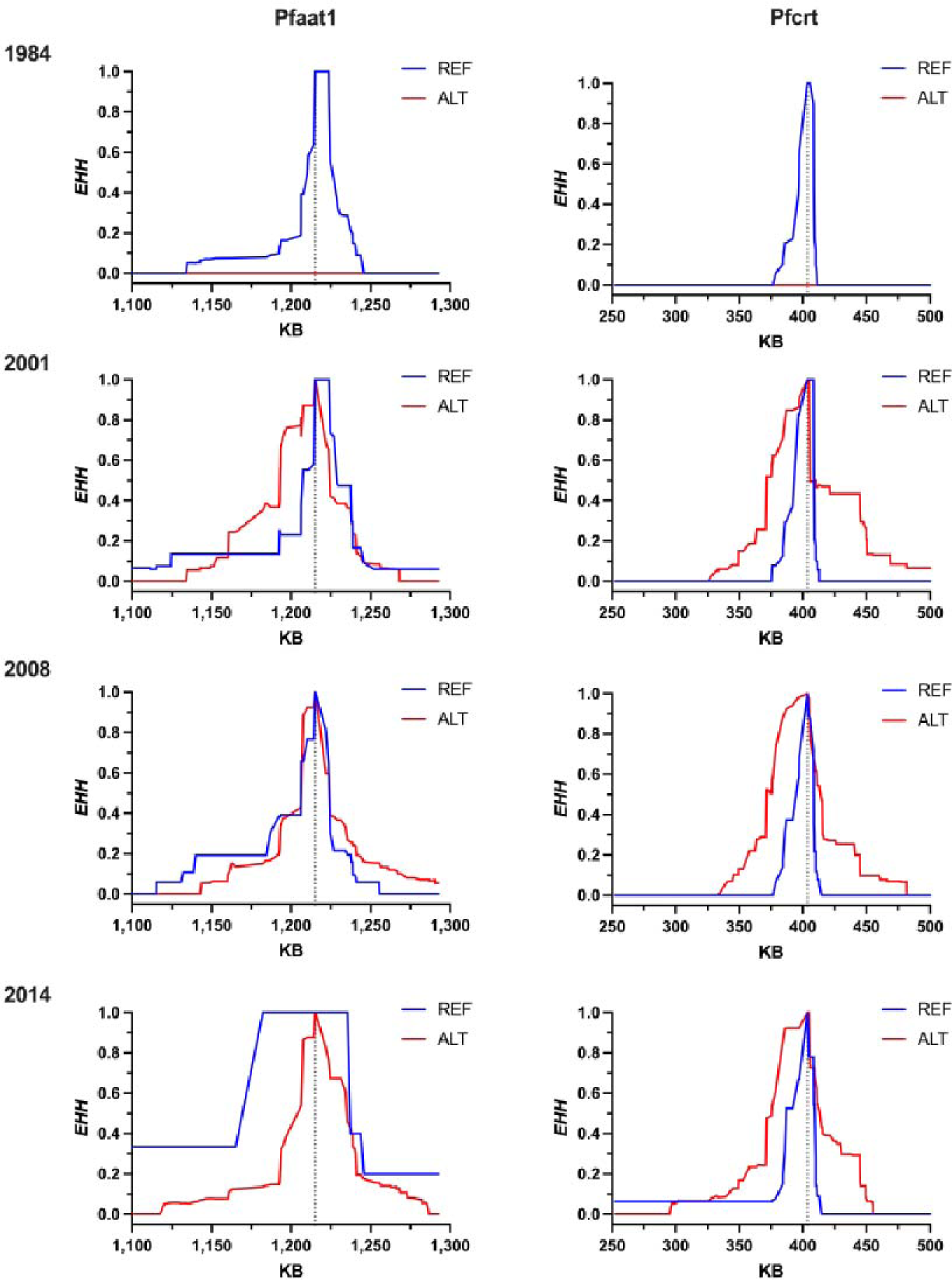
Extended haplotype homozygosity (EHH) analysis of Gambian samples. Graphs show EHH surrounding *pfaat1* S258<u>L</u> (left panels), or *pfcrt* K76<u>T</u> (right panels) in samples collected in 1984, 2001, 2008 and 2014. 1990 was omitted because only 13 samples were available. Sample from 1984 show EHH around the ancestral allele only (blue), because neither *pfaat1* S258<u>L</u> nor *pfcrt* K76<u>T</u> were sampled at that time.

**Fig. S6.**
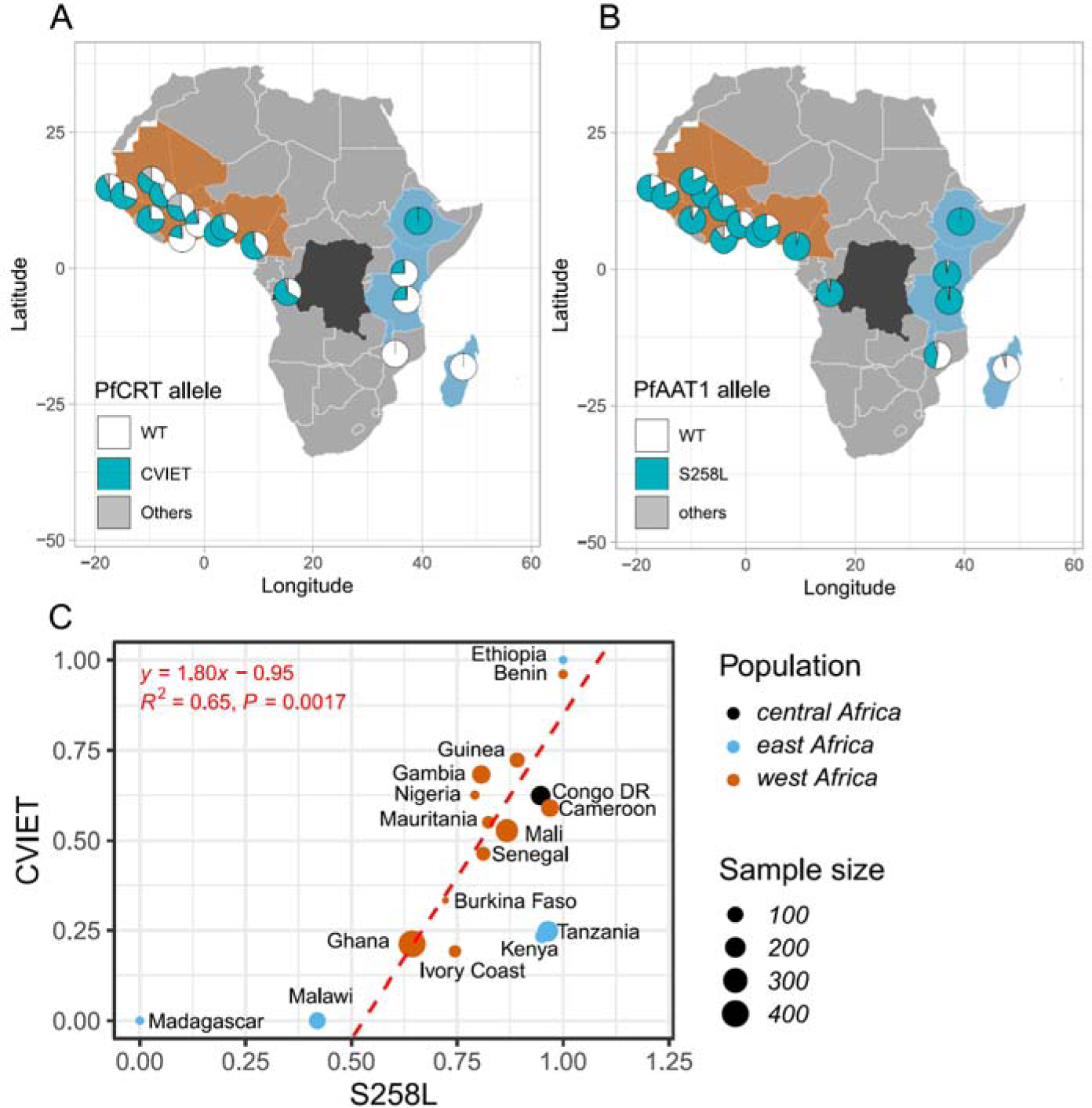
*pfcrt* and *pfaat1* allele frequency distributions and correlations in African countries. A. *pfcrt* allele distribution in African countries. B. *pfaat1* allele distribution in African countries. C. Correlations in allele frequencies between *pfcrt* (CVIET) and *pfaat1* (S258<u>L</u>). Frequencies of the CVIET haplotype for amino acids 72-76 in *pfcrt* are significantly correlated with allele frequencies of *pfaat1* S258<u>L</u> in West Africa (*R^2^* = 0.65, *p* = 0.0017, red dashed line) or across all African populations (*R^2^* = 0.44, *p* = 0.0021). Point size indicates sample numbers, while color indicates sampling locations.

**Fig. S7.**
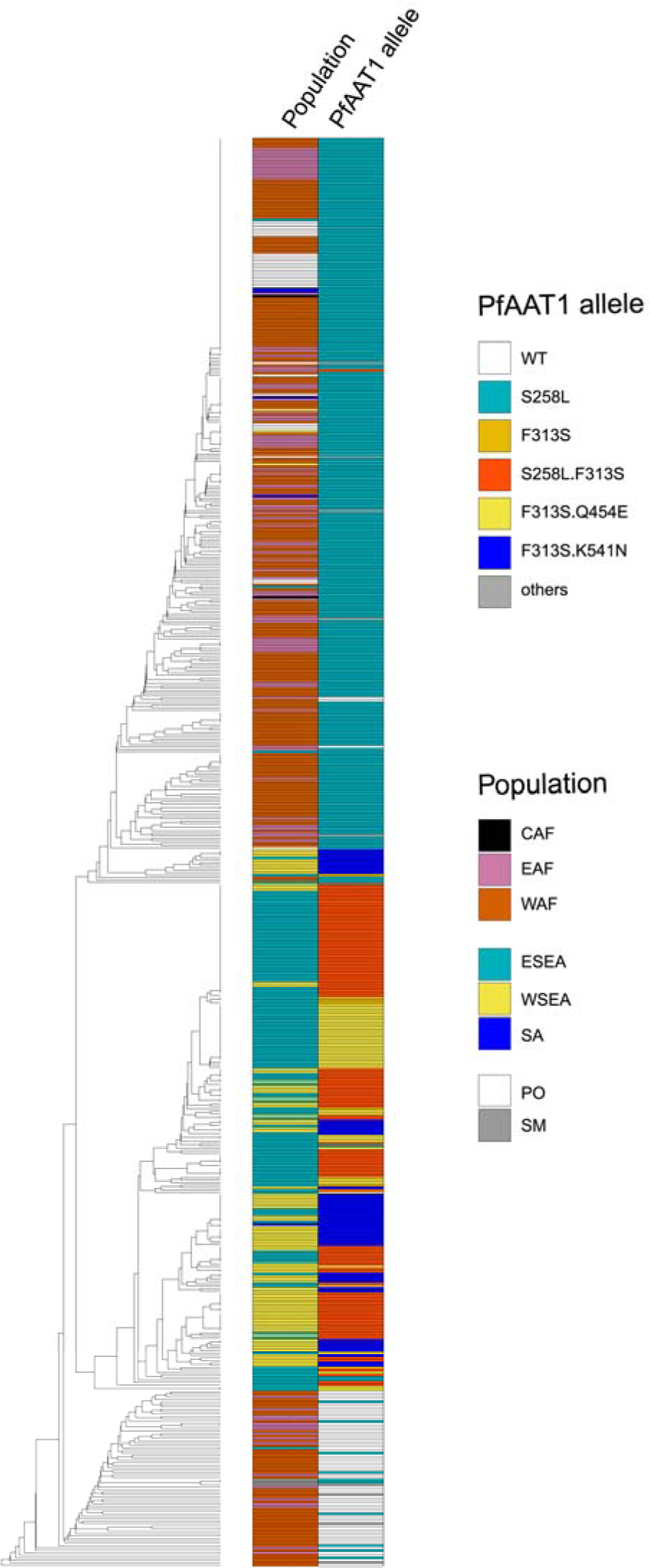
UPGMA tree showing the relationship of 581 haplotypes based on SNPs inside the 50 kb region surrounding *pfaat1*. The tree was rooted with *Plasmodium reichenowi* (not shown in the tree). WAF: west Africa, EAF: east Africa, CAF: central Africa, SM: south America, ESEA: east Southeast (SE) Asia, SA: south Asia, WSEA: west SE Asia, PO: Pacific Ocean.

**Fig. S8.**
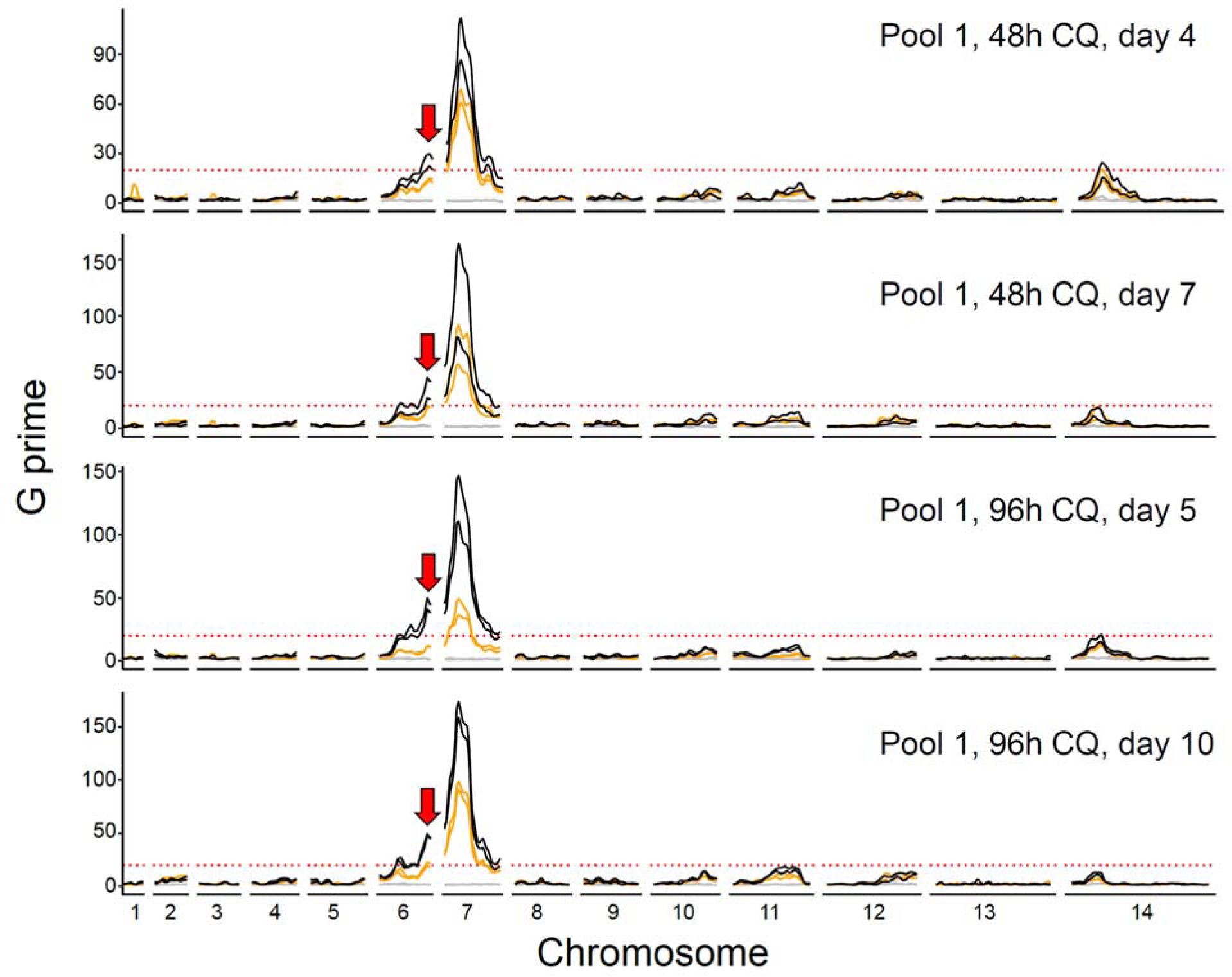
Mapping CQ resistance loci with cross 3D7×NHP4026 using recombinant progeny pool 1. Grey, orange, and black lines indicate BSAs from 50 nM, 100 nM, and 250 nM CQ treatments, separately. Lines with the same color are BSAs from technical replicates. Red dashed lines are the threshold (*G prime* = 20) for QTL detection. Red arrows indicate the location of the chromosome 6 QTL.

**Fig. S9.**
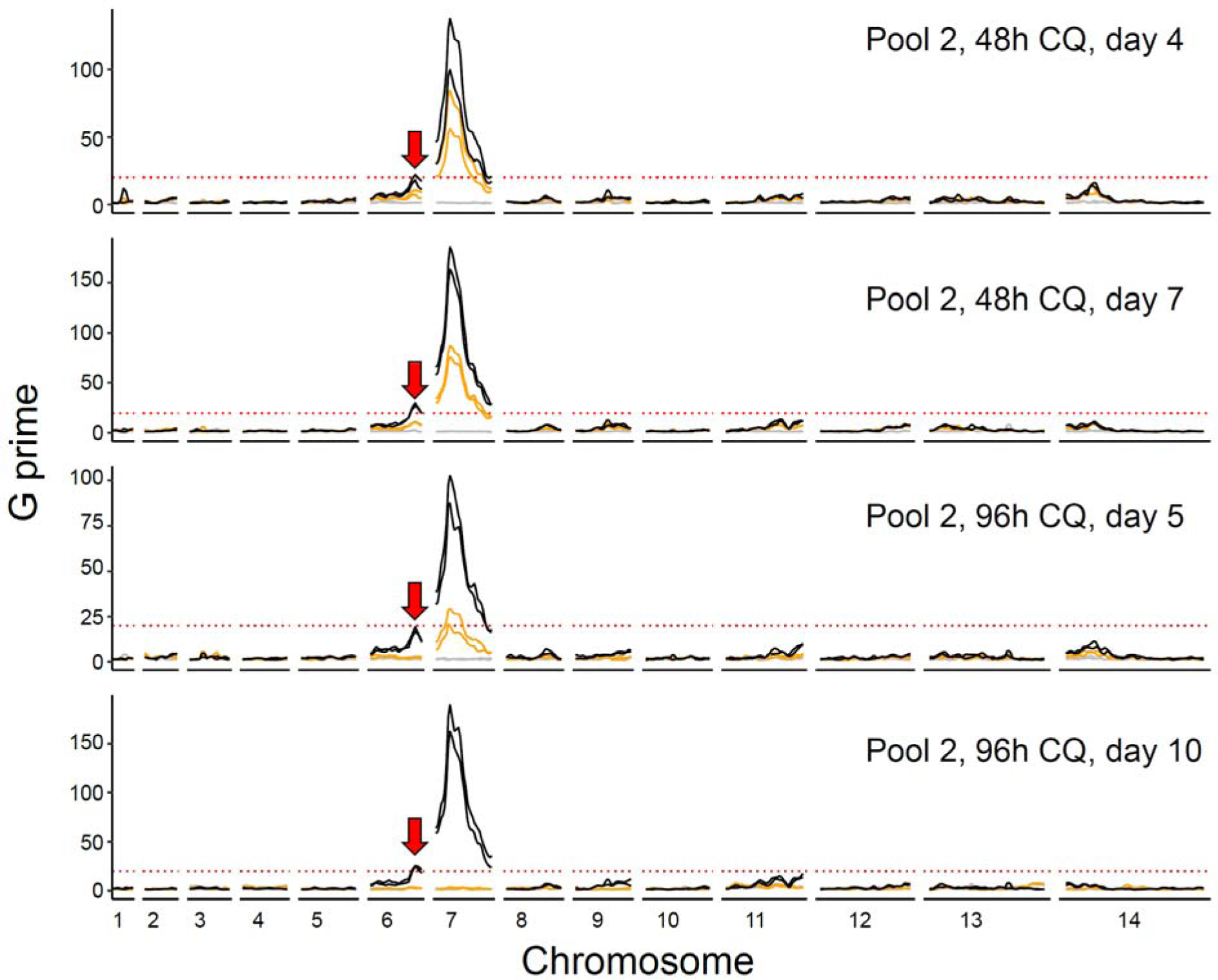
Mapping CQ resistance loci with cross 3D7×NHP4026 using recombinant progeny pool 2. Grey, orange, and black lines indicate BSAs from 50 nM, 100 nM, and 250 nM CQ treatments, separately. Lines with the same color are BSAs from technical replicates. Red dashed lines are the threshold (*G prime* = 20) for QTL detection. Red arrows indicate the location of the chromosome 6 QTL.

**Fig. S10.**
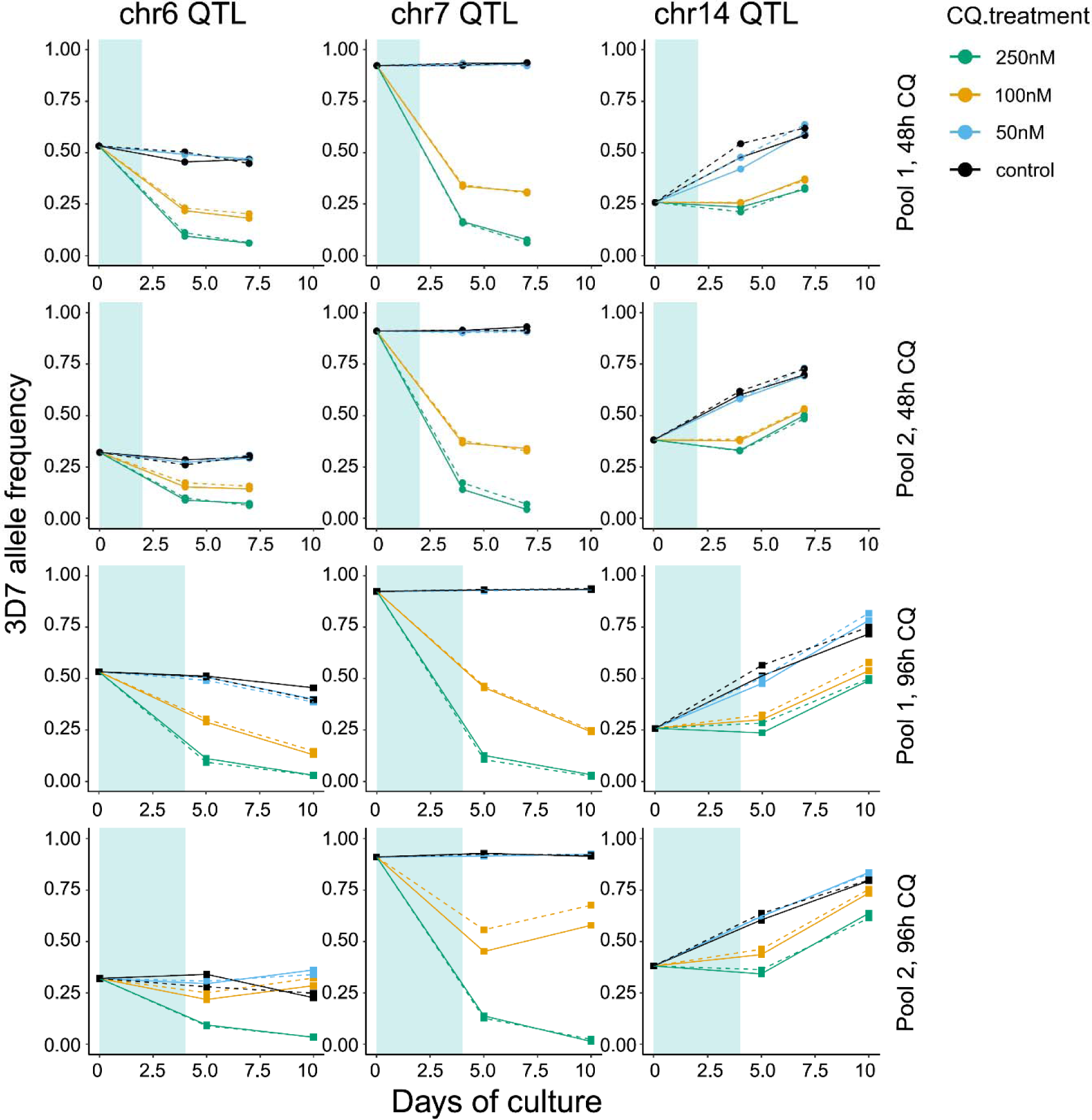
3D7 allele frequency for QTLs at chr.6, chr.7 and chr.14. CQ treatments were applied to the pools on day 0 at 0 (control), 50, 100, or 250 nM. CQ was removed on day 2 (48 hour treatment) or day 4 (96 hour treatment), as shaded with light blue. For 48 hour CQ treatments, samples were collected at day 0, 4, and 7; while for 96 hour CQ treatment, samples were collected at day 0, 5, and 10. Solid or dashed lines are results from different technical replicates. The 3D7 allele frequencies in CQ treated pools decrease at both chr.6 and chr.7 QTL regions. At the chr.14 QTL region, allele frequencies are unchanged following drug treatment, suggesting this QTL is unrelated to drug treatment.

**Fig. S11.**
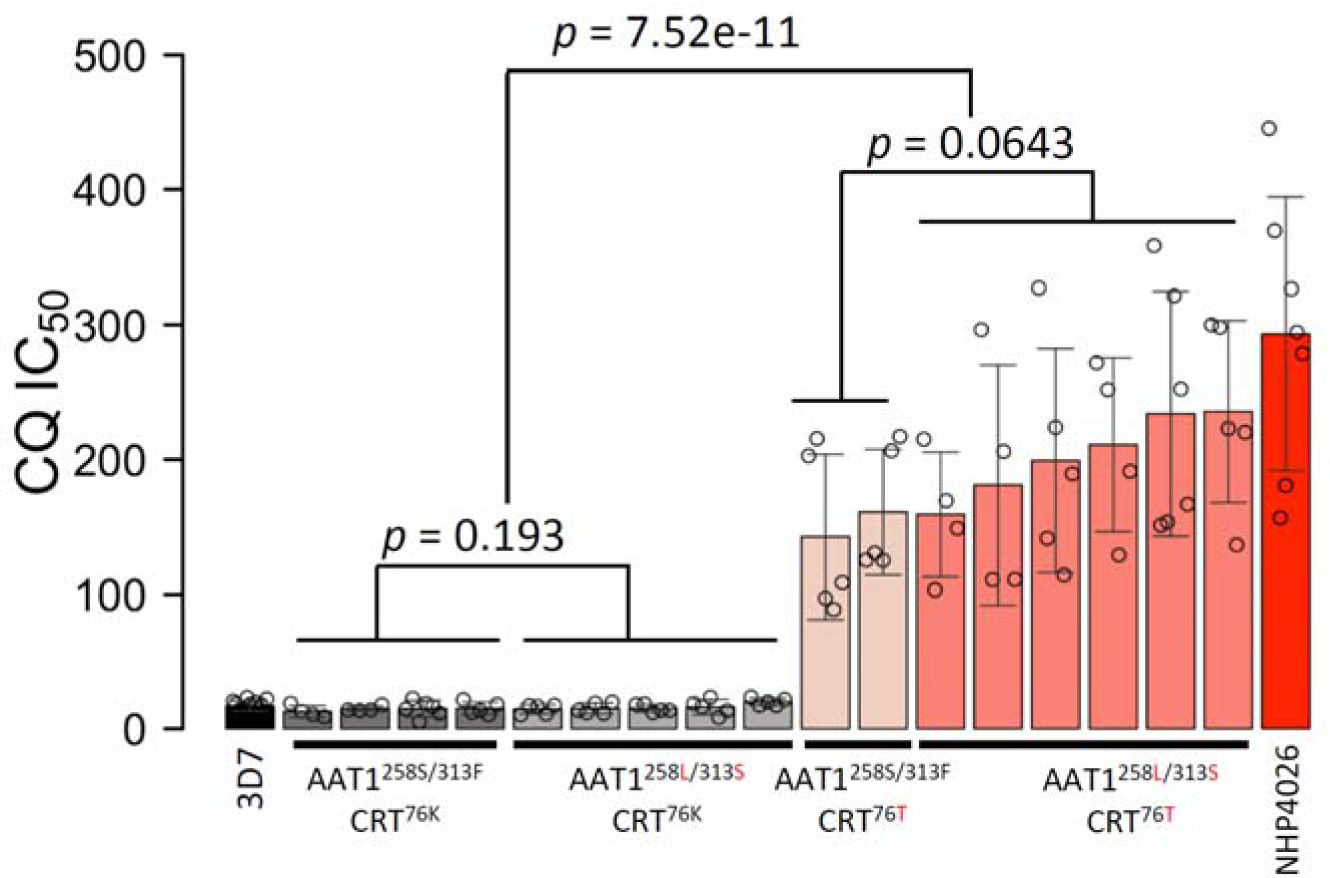
Mean IC_50_ (± S.D.) of parents and progeny grouped by combinations of *pfcrt* and *pfaat1* allele. IC_50_ for each clone is calculated from 4-13 biological replicates (Supplementary Table 7). Only two progeny carrying *pfaat1* 258S/313F (WT) and *pfcrt* K76<u>T</u> were recovered. *P* values indicate significance levels are based on two-way ANOVA analysis.

**Fig. S12.**
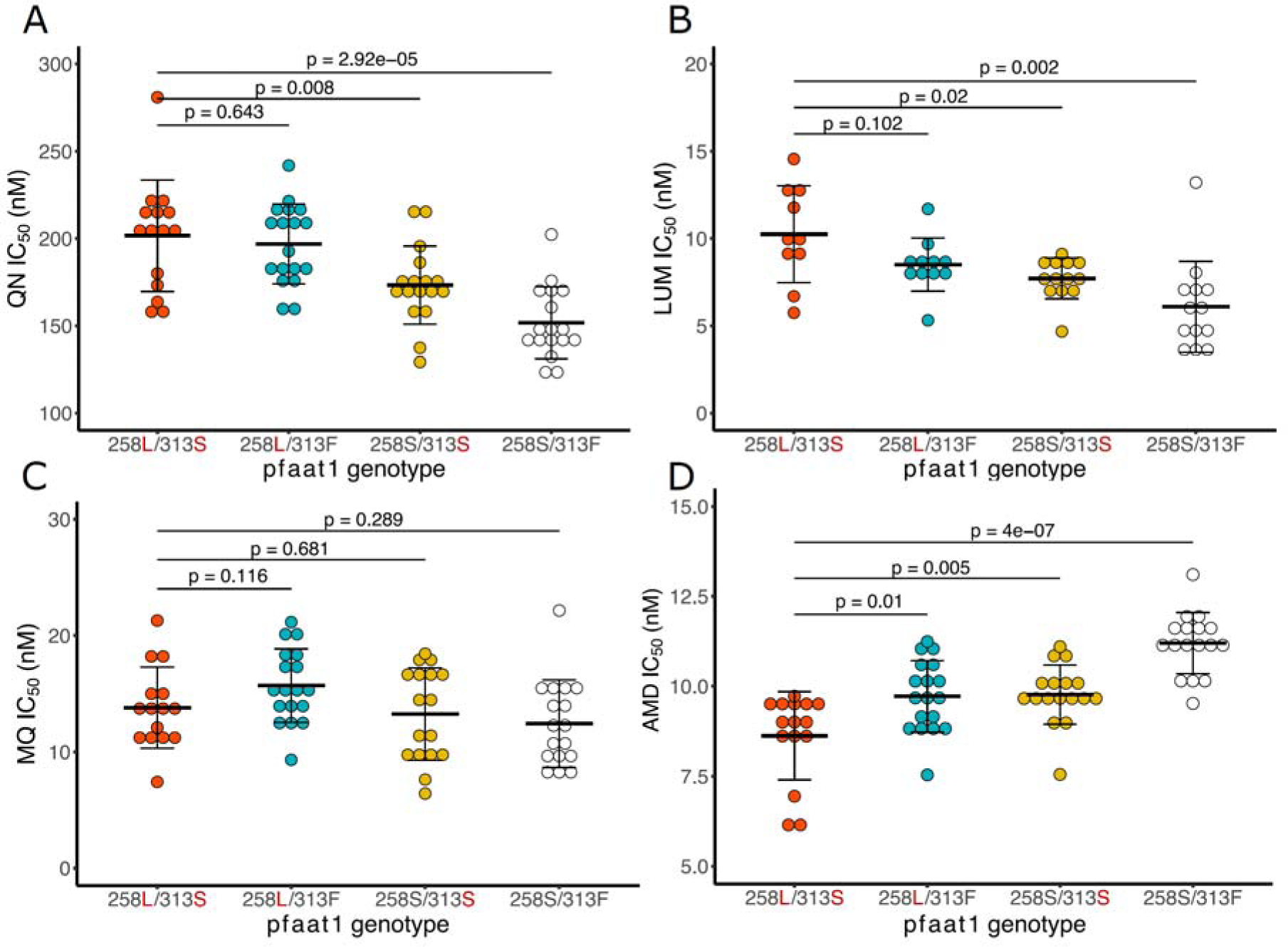
Impact of CRISPR/Cas9 substitutions on IC_50_ of quinine, lumefantrine, mefloquine and amodiaquine. CRISPR/Cas9 gene editing resulted in small differences in IC_50_ for quinine (QN), lumefantrine (LUM) and Amodiaquine (AMD), but no significant changes for mefloquine (MQ). However, all IC_50’_s were below levels of clinical significance for these drugs (clinical thresholds: QN = 600 nM ^67^; MQ = 30 nM ^67^, AMD = 60 nM ^67^), or at the lower end of the *in vitro* range (0-150 nM) in the case of LUM ^68^. *P* values indicate significance levels are based on two-way ANOVA analysis.

**Fig. S13.**
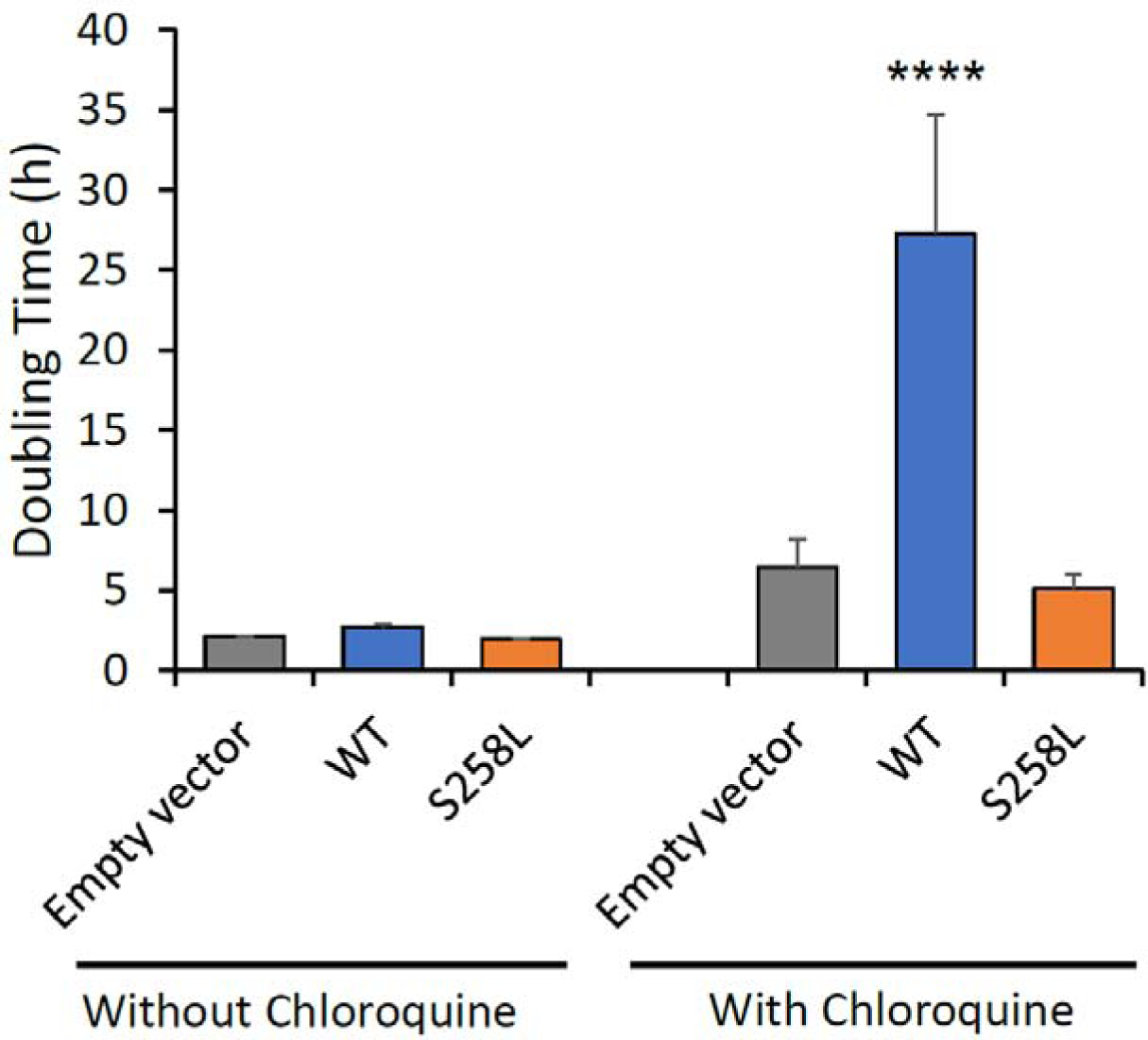
Introduction of S258<u>L</u> results in chloroquine resistance in yeast. Yeast doubling time was calculated from the linear portion of exponential growth. Data was shown as means from 3 independent experiments ± SEM, and significance and was calculated according to multiple comparisons (with Turkey corrections) of two-way ANOVA. Growth of yeast cells expressing wild type *pfaat1* (WT) is severely impacted by CQ treatment (5 mM CQ through the experiments) but is recovered in yeast expressing *pfaat1* S258<u>L</u>. Published results demonstrate that AAT1 is expressed in the yeast cell membrane ^26^.

**Fig. S14.**
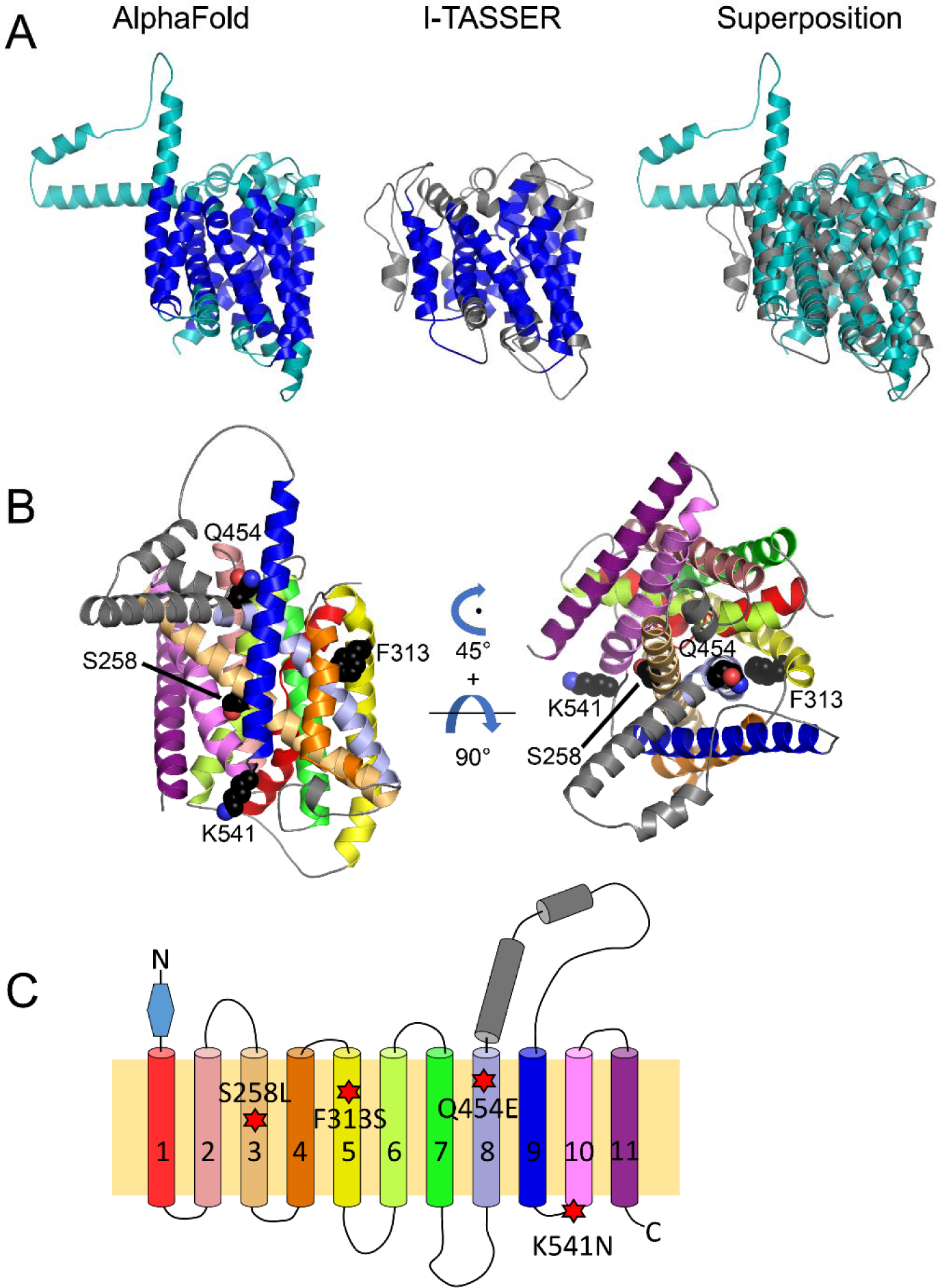
Topology structure of pfAAT1 protein. (A) AlphaFold model of PfAAT1 (left), representative I-TASSER model of PfAAT1 (center), structural superposition of the AlphaFold model (teal) and I-TASSER model (gray, right). TOPCONS transmembrane (TM) helix topology predictions are mapped onto the models in dark blue (left, center). AlphaFold and I-TASSER models align with a RMSD of 2.5 Å over 327 of 440 residues. Amino-terminal residues 1-166 were excluded from all models due to low confidence in structure prediction. (B) Detailed view of the mutations on the predicted PfAAT1 3-D structure using the AlphaFold model. The right view is related to the left by a 45° rotation about the axis looking down at the figure followed by a 90° rotation about the horizontal axis. The four SNPs shown as space-filling models are all arranged within a plane at one side of the model, perpendicular to the membrane. S258<u>L</u> (helix 3) and F313<u>S</u> (helix 5) are located opposite each other with helix 8 in between. Given the epistatic interactions between the PfAAT1 S258<u>L</u> and F313<u>S</u> SNPs evident from our functional analyses, the F313<u>S</u> substitution of the bulky, hydrophobic phenylalanine with the smaller, polar serine may compensate for a disruption in the transmembrane region that includes helices 3, 5, and 8 allowing for partial restoration of amino acid transport activity. Q454<u>E</u> is located on helix 8 near the TM surface and K541<u>N</u> is located in a loop connecting helix 9 and 10. (C) Topology of PfAAT1 inferred using 3D structure. There are eleven transmembrane (TM) helices. Three of the mutations are located at the TM helices, while K541<u>N</u> is located at a loop connecting helix 9 and 10. The color scheme matches the schematic in Panel B. The blue triangle indicates amino-terminal residues 1-166 that were excluded from structure prediction.

**Fig. S15.**
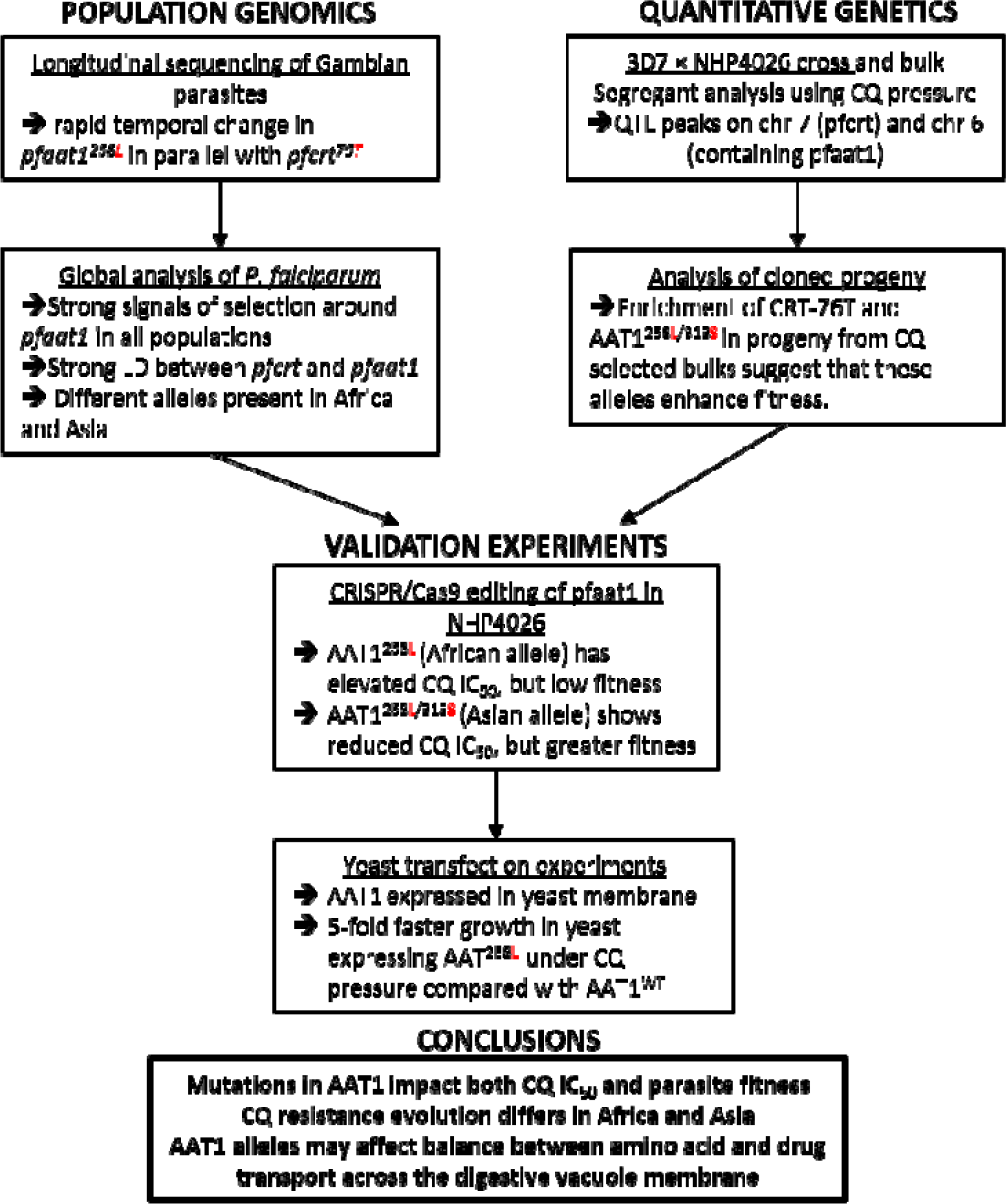
Project design. We use (i) population genomic analyses, (ii) genetic crosses and quantitative genetics analysis followed by (iii) functional analyses to investigate the role of additional loci in CQ resistance.

**Fig. S16.**
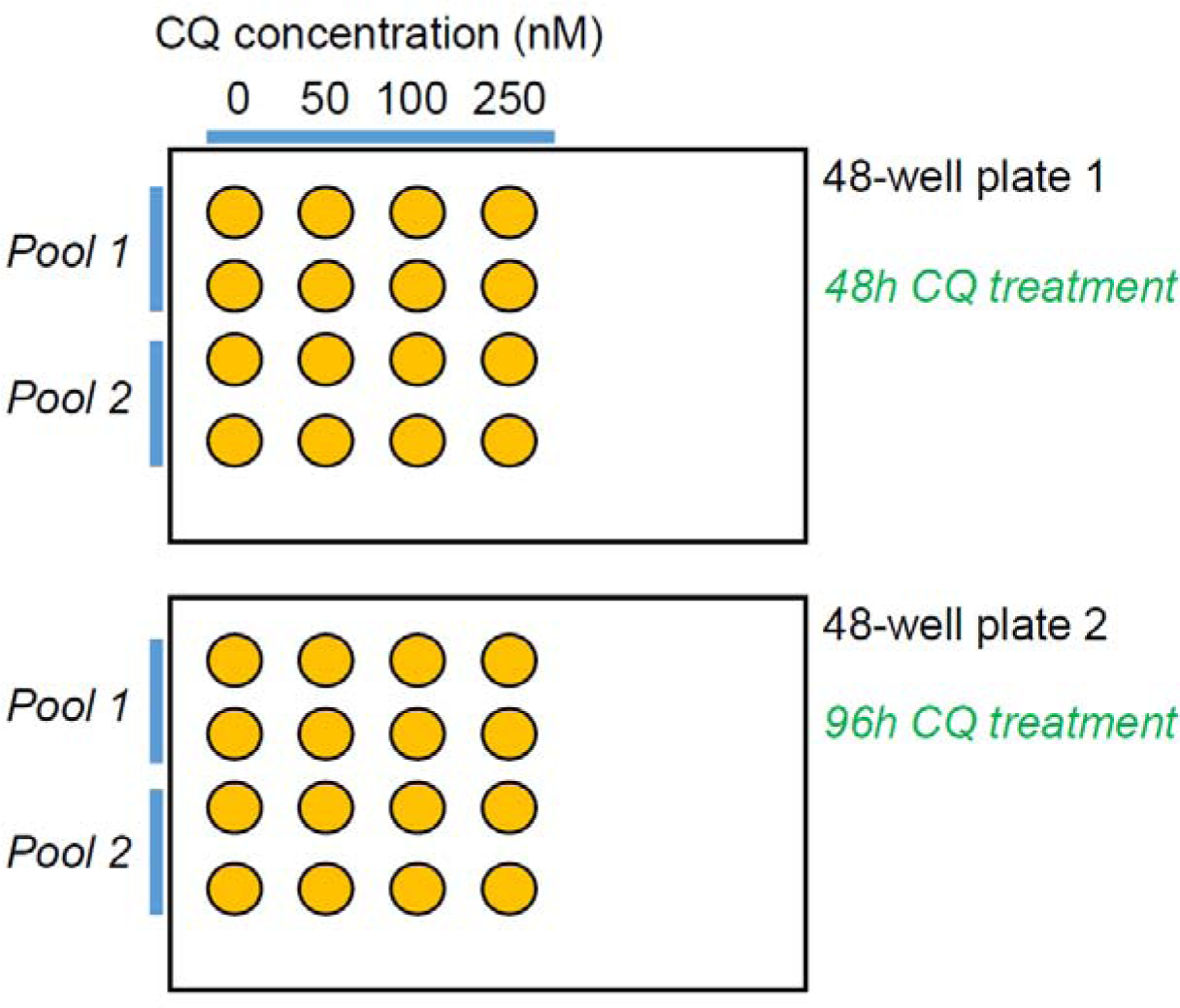
Experiment design for CQ bulk segregant analysis.

**Fig. S17.**
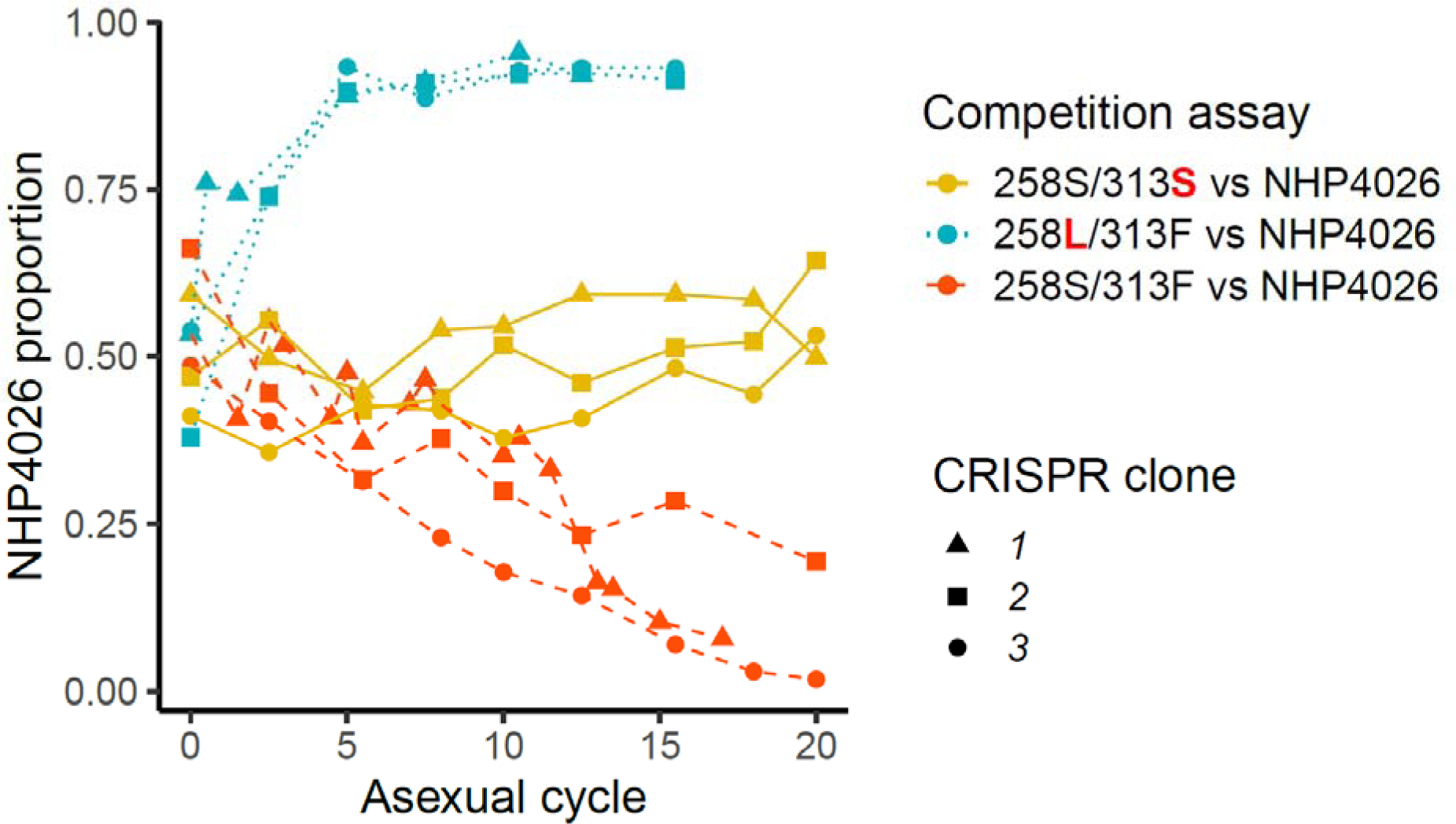
Allele frequency changes in head-to-head competition experiments between different CRISPR/Cas9 edited parasites and NHP4026. We used three independent CRISPR/Cas9 edited clones for each genotype, and two technical replicates for each competition experiment (average values plotted).

## Notes

### Competing Interest Statement

The authors have declared no competing interest.

### Summary of Updates

Significant changes have been made to the article in response to peer review. These changes include: additional population genetic analysis including haplotype analysis of pfCRT, correlation of AAT-S258L and CRT-K76T across African locations, further analysis of LD between AAT1-S258L and CRT-K76T in field populations, new analysis of IBD by geographical region, reanalysis of IC50 data from progeny clones and analysis of pfAAT-haplotypes inherited in all cloned progeny.

